# EZH2 inhibition remodels the inflammatory senescence-associated secretory phenotype to potentiate pancreatic cancer immune surveillance

**DOI:** 10.1101/2022.06.21.495523

**Authors:** Loretah Chibaya, Katherine C. Murphy, Yvette Lopez-Diaz, Kelly D. DeMarco, Haibo Liu, Sneha Gopalan, Melissa Faulkner, Junhui Li, John P. Morris, Yu-jui Ho, Janelle Simon, Wei Luan, Amanda Kulick, Elisa de Stanchina, Karl Simin, Lihua Julie Zhu, Thomas G. Fazzio, Scott W. Lowe, Marcus Ruscetti

## Abstract

T cell-activating immunotherapies that produce durable and even curative responses in some malignancies have failed in pancreatic ductal adenocarcinoma (PDAC) due to rampant immune suppression and poor tumor immunogenicity. We and others have demonstrated that induction of cellular senescence and its accompanying senescence-associated secretory phenotype (SASP) can be an effective approach to activate not only T cell but also cytotoxic Natural Killer (NK) cell-mediated anti-tumor immunity. Here we found that the pancreas tumor microenvironment (TME) suppresses NK and T cell surveillance following therapy-induced senescence through EZH2-mediated repression of pro-inflammatory SASP genes. Genetic or pharmacological inhibition of EZH2 or its methyltransferase activity stimulated the production of pro-inflammatory SASP chemokines CCL2 and CXCL9/10 that led to enhanced NK and T cell infiltration and tumor eradication in preclinical PDAC mouse models. EZH2 activity was also associated with suppression of SASP-associated inflammatory chemokines and cytotoxic lymphocyte immunity and reduced overall survival in a PDAC patient cohort. These results demonstrate that EZH2 mediates epigenetic repression of the pro-inflammatory SASP in the pancreas TME, and that EZH2 blockade in combination with senescence-inducing therapies could be a powerful means to potentiate NK and T cell surveillance in PDAC to achieve immune-mediated tumor control.

## INTRODUCTION

Pancreatic ductal adenocarcinoma (PDAC) is a devastating disease with few effective treatment options, and has quickly risen to become the 3^rd^ leading cause of cancer-related death^1^. Conventional chemotherapy regimens have limited efficacy in PDAC, in part due to a fibrotic and desmoplastic tumor microenvironment (TME) that leads to vascular dysfunction and poor drug delivery and activity in tumors^2–4^. Immunotherapy regimens including chimeric antigen receptor (CAR) T cells and anti-PD-1 and CTLA-4 immune checkpoint blockade (ICB) therapies that have been effective in other aggressive, chemo-refractory tumors have been ineffective in PDAC because of widespread innate and adaptive immune suppression in the pancreas TME^5–7^. Indeed, an abundance of suppressive macrophage and myeloid populations, poor tumor immunogenicity, and a lack of cytotoxic Natural Killer (NK) and T cell infiltration contribute to the immunologically “cold” TME associated with PDAC and immunotherapy resistance^8^. Thus, new and innovative approaches are needed to target the multiple axes of immune suppression in PDAC to achieve durable therapeutic outcomes.

Point mutations in KRAS are oncogenic drivers in PDAC and found in >90% of patients^9^. One strategy to increase tumor immunogenicity and stimulate anti-tumor immunity is to target the oncogenic pathways that drive immune suppression^10,11^, including RAS signaling itself^12,13^. RAS pathway targeting therapies have been shown not only to increase tumor immunogenicity through upregulating antigen presentation and processing genes (e.g. major histocompatibility complex (MHC) Class I (MHC-I) molecules) but also lead to immune stimulatory microenvironments that activate anti-tumor NK and T cell immunity and ICB therapy efficacy^14–18^. We previously showed that the combination of the MEK inhibitor trametinib and CDK4/6 inhibitor palbociclib (T/P) could trigger KRAS mutant cancers to enter cellular senescence, a stable cell cycle arrest program that is accompanied by a secretory program that can modulate immune responses^19,20^. This senescence-associated secretory phenotype (SASP) includes a collection of pleiotropic factors such as pro- and anti-inflammatory chemokines and cytokines, angiogenic factors, growth and stemness components, matrix metalloproteinases (MMPs), and lipid species that remodel the surrounding TME in both tumor promoting and tumor suppressive ways depending on the context^21–23^.

In certain tumor and cancer therapy contexts, we and others have demonstrated that the SASP can mediate potent anti-tumor immunity to block tumor formation, regress established tumors, and enhance immunotherapy regimens^19,20,24–27^. Recently we found that therapy-induced senescence following T/P treatment could induce anti-tumor immune surveillance in preclinical mouse models of KRAS mutant lung adenocarcinoma (LUAD) and PDAC. In KRAS mutant LUAD, T/P-induced senescence led to secretion of pro-inflammatory SASP factors that activated NK cell immune surveillance and drove NK cell-mediated long-term lung tumor responses^19^. Intriguingly, in similar genetic models of KRAS mutant PDAC, T/P treatment led to a predominantly pro-angiogenic SASP that enhanced vascularization and CD8^+^ T cell extravasation into PDAC with little effect on NK cell immunity^20^. Combining therapy-induced senescence with anti-PD-1 ICB enhanced cytotoxic T cell immunity and led to tumor responses in PDAC-bearing animals, demonstrating that the SASP could be a means to make “cold” PDAC tumors “hot” and potentiate currently ineffective ICB strategies.

In order to effectively harness senescence and its immune stimulating properties for tumor suppression, we need a better understanding of why the SASP elicits altered immune responses in different cancer contexts and how the SASP transcriptional program or specific SASP factors can be optimized for immune-mediated tumor destruction^28^. In the setting of PDAC, it will be important to elucidate how the pancreas TME suppresses cytotoxic NK cells that can act as potent eliminators of senescent tumor cells^29^. As NK cells can both directly eradicate target cells through release of cytolytic granules, as well as indirectly mobilize adaptive T cell immunity through secretion of cytokines and chemokines, they are promising targets for cancer immunotherapy^30^. Here we set out to address why the SASP elicited different immune responses in the pancreas and how it could be harnessed for NK cell immunotherapy in PDAC.

## RESULTS

### NK cell immunity is induced in the lung but not pancreas TME following therapy-induced senescence

Based on our previous findings we hypothesized that the pancreas TME may contribute to suppression of NK cell anti-tumor immunity following therapy-induced senescence. To test this, we took advantage of genetically similar KRAS mutant PDAC and LUAD cell lines that could be transplanted syngeneically into different organs of C57BL/6 mice to study the impact of the TME on senescence-driven immune responses. These included: (a) *KPC* PDAC tumor cell lines (*KPC1*, *KPC2*) derived from PDAC tumors in *Pdx1-Cre*; *LSL-KRAS^G12D^*;*Trp53^R172H/wt^*genetically engineered mouse models (GEMMs)^31^ and (b) *KP* LUAD cell lines (*KP1*, *KP2*) derived from lung tumors in *Kras^LSL-G12D/wt^;Trp53^flox/flox^* GEMM mice administered an adenovirus expressing Cre-recombinase intratracheally^32^. *KPC* PDAC or *KP* LUAD cells were engineered to express luciferase and GFP (to track tumors *in vivo*) and then transplanted intravenously (i.v.) to form tumors in the lungs or injected directly into the pancreas of C57BL/6 mice (Fig. 1a,b). Additionally, PDAC and LUAD cells were also transplanted into the liver, a common site of metastasis for both tumor types (Fig. 1c). Following tumor formation, mice were treated with vehicle or T/P for two weeks to induce senescence (Fig. 1a-c). PDAC and LUAD tumors propagated in each organ had a similar tumor burden and disease histopathology, as well as senescence (senescence-associated β-galactosidase (SA-β-gal)), anti-proliferative (Ki67), and on-target drug responses (pERK, pRb) to T/P treatment (Extended Data Fig. 1a,b).

**Fig. 1.**
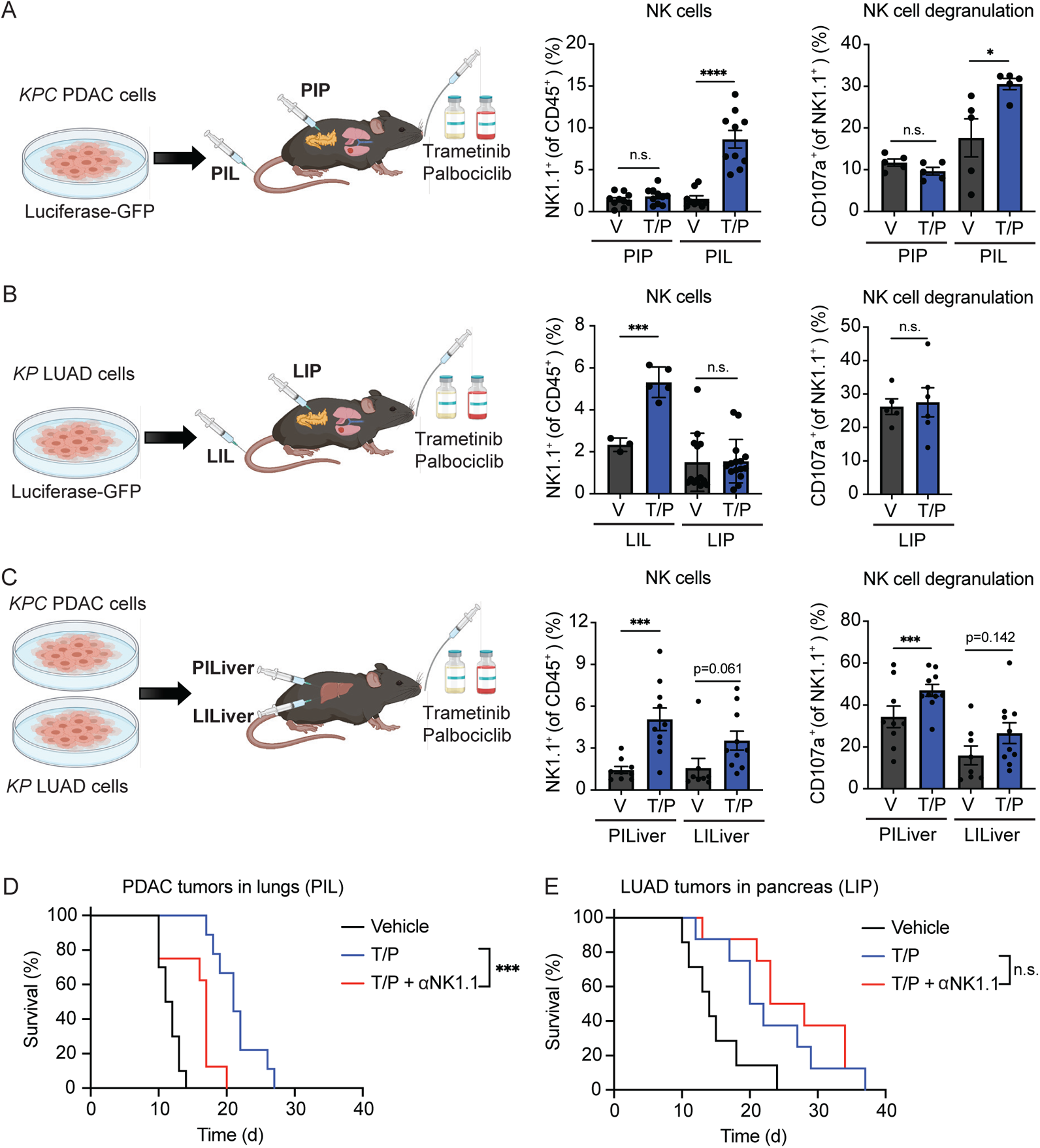
NK cell immunity is induced in the lung but not pancreas TME following therapy-induced senescence. **a-b**, *KPC1* PDAC (**a**) or *KP1* LUAD (**b**) tumor cells expressing luciferase-GFP were injected i.v. or orthotopically into the pancreas of C57BL/6 mice. Following tumor formation in the lungs or pancreas, mice were treated with vehicle (V) or combined trametinib (1mg/kg body weight) and palbociclib (100 mg/kg body weight) (T/P) for 2 weeks (left). Right, flow cytometry analysis of NK cell numbers and degranulation in each condition (n ≥ 3 per group). **c**, *KPC1* PDAC or *KP1* LUAD cells expressing luciferase-GFP were injected orthotopically into the livers of C57BL/6 mice and treated as in (**a**) following tumor formation (left). Right, flow cytometry analysis of NK cell numbers and degranulation in each condition following 2-week treatment (n ≥ 8 per group). **d**, Kaplan-Meier survival curve of C57BL/6 mice harboring *KPC1* PDAC tumors in lungs (PIL) treated with vehicle or trametinib (1 mg/kg) and palbociclib (100 mg/kg) in the presence or absence of a NK1.1 depleting antibody (PK136; 250 ug) (n ≥ 8 per group). **e**, Kaplan-Meier survival curve of C57BL/6 mice harboring *KP1* LUAD tumors in pancreas (LIP) and treated as in (**d**) (n ≥ 7 per group). PIP, PDAC tumors in pancreas; PIL, PDAC tumors in lungs; LIL, LUAD tumors in lungs; LIP, LUAD tumors in pancreas. PILiver, PDAC tumors in liver; LILiver, LUAD tumors in liver. *P* values in **a-c** were calculated using two-tailed, unpaired Student’s t-test, and those in **d** and **e** calculated using log-rank test. Error bars, mean ± SEM. **** *P* <0.0001, *** *P* <0.001, * *P* <0.05. n.s., not significant.

In line with our previous findings, T/P treatment led to increased NK cell accumulation and cytotoxicity, as marked by the degranulation markers CD107a and Granzyme B (GZMB), in LUAD tumors grown in the lungs (LIL) but not in PDAC tumors grown in the pancreas (PIP) despite peripheral NK cell expansion in adjacent spleens (Fig. 1a,b and Extended Data Fig. 1c-e). NK cell suppression was specific to the pancreas TME, as PDAC tumors grown in the lungs (PIL) or liver (PILiver) underwent NK cell surveillance following T/P-induced senescence (Fig. 1a,c and Extended Data Fig. 1c). Similarly, whereas LUAD tumors propagated in the liver (LILiver) were infiltrated with activated NK cells, those propagated in the pancreas (LIP) were not (Fig. 1b,c and Extended Data Fig. 1d). These tissue-specific changes in NK cell states were functionally relevant, as NK cell depletion with an NK1.1-targeting antibody (PK136) reduced the survival benefit of T/P treatment in mice bearing tumors in the lungs (PIL) but not those with tumors in the pancreas (LIP, PIP) (Fig. 1d,e and Extended Data Fig. 1f). In contrast, T/P-induced senescence led to increased CD4^+^ and CD8^+^ T cells in all tumor-bearing tissues, though the infiltrating T cells were not activated and T cell depletion studies indicated they did not contribute to anti-tumor immunity in the lung or pancreas TME (Extended Data Fig. 1f-h). Therefore, the pancreas TME leads to specific resistance to NK cell immune surveillance following therapy-induced senescence.

### The pancreas TME suppresses the pro-inflammatory SASP

We next performed RNA-sequencing (RNA-seq) on GFP-labeled PDAC and LUAD cells FACS sorted from lung or pancreas tumors following T/P treatment to determine the impact of the TME on signaling in senescent tumor cells (Fig. 2a). T/P treatment led to significant enrichment of inflammatory pathways related to NF-κB, TNF, and chemokine signaling, as well as type I interferon and IL-12 pathways known to activate innate and in particular NK cell immunity, in PDAC and LUAD cells in the lungs (PIL, LIL) as compared to those in the pancreas (LIP, PIP) (Fig. 2b, Extended Data Fig. 2a, and Supplementary Table 1). A subset of pro-inflammatory SASP genes were significantly upregulated following T/P-induced senescence in PDAC and LUAD cells in the lung TME but not those in the pancreas TME (Fig. 2c). These included a number of SASP-related chemokines known to regulate the chemotaxis of NK cells and T cells into tumors, including CCL2, CCL5, CCL7, CCL8, CXCL9, and CXCL10 (Fig. 2d)^33^. This pancreas tissue-specific suppression of the SASP output was independent of senescence-associated cell cycle arrest, as T/P treatment uniformly reduced E2F and MYC target gene expression, the Ki67 proliferation index, and other senescence-related markers and gene sets in all tumor conditions tested (Extended Data Fig. 1b and 2b,c). *In vitro*, T/P treatment induced the expression and secretion of pro-inflammatory SASP factors to a similar degree in *KPC* PDAC and *KP* LUAD cells and in KRAS mutant human PDAC and LUAD cell lines (Extended Data Fig. 3), suggesting that PDAC cells have the intrinsic capacity to produce pro-inflammatory SASP factors following therapy-induced senescence. Thus, a set of pro-inflammatory SASP genes known to regulate NK cell immune surveillance are transcriptionally repressed in the pancreas TME following therapy-induced senescence.

**Fig. 2.**
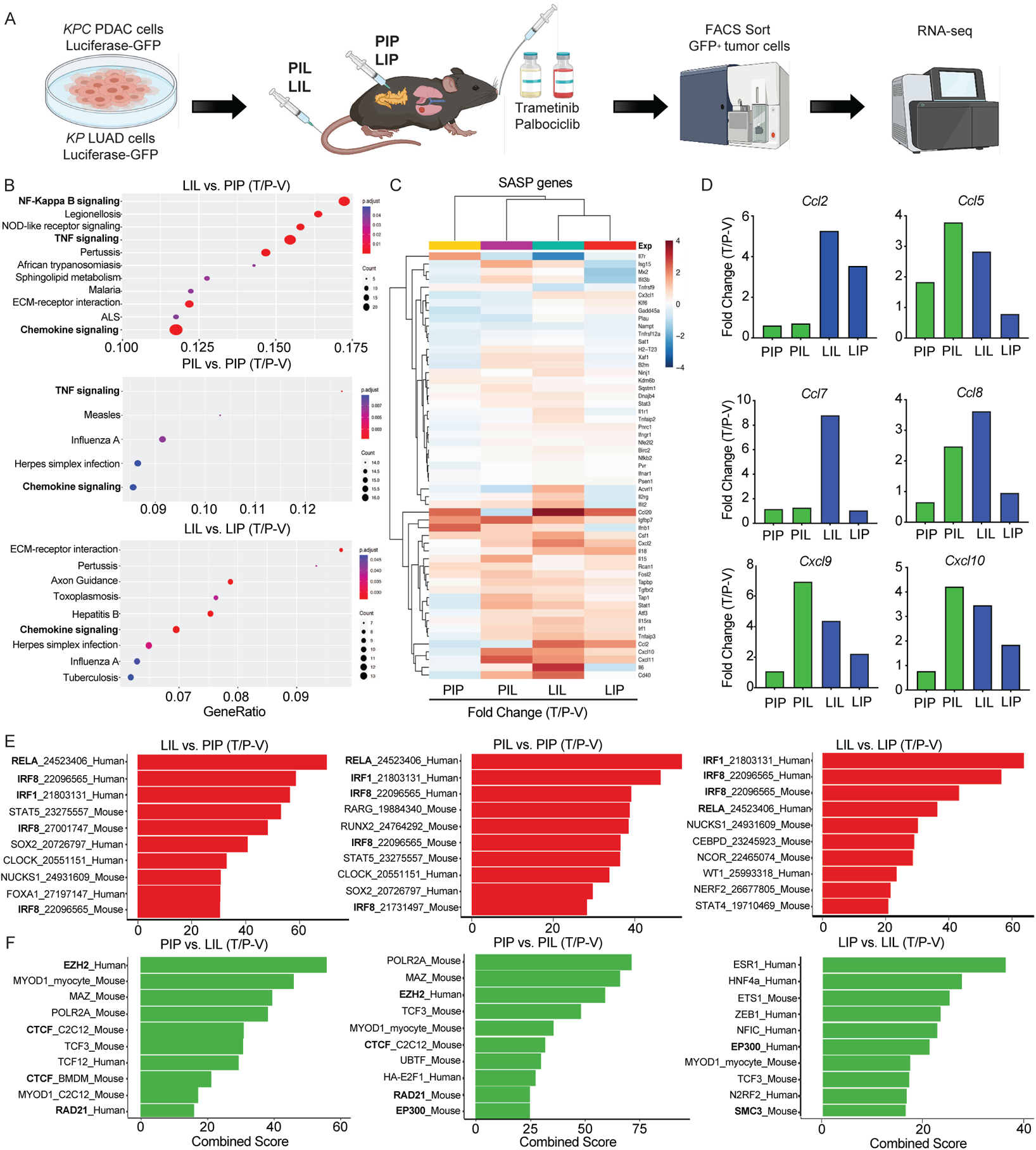
The pancreas TME suppresses the pro-inflammatory SASP. **a**, *KPC1* PDAC or *KP1* LUAD tumor cells expressing luciferase-GFP were injected i.v. or orthotopically into the pancreas of C57BL/6 mice. Following tumor formation in the lungs or pancreas, mice were treated with vehicle (V) or combined trametinib (1mg/kg) and palbociclib (100 mg/kg) (T/P) for 2 weeks (left). GFP^+^ tumor cells were FACS sorted and extracted RNA subjected to RNA-seq analysis (n=2-4 per group). **b**, KEGG pathway analysis of pathways enriched in tumors in the lungs (LIL, PIL) compared to tumors in the pancreas (PIP, LIP) following T/P treatment. **c**, Heatmap showing fold change in SASP gene expression following T/P treatment in indicated tumor settings. **d**, Fold change in expression of select SASP chemokines following T/P treatment in indicated tumor settings. **e-f,** Transcription factor enrichment analysis showing transcriptional regulators whose targets are enriched in tumors in the lungs (LIL, PIL) (**e**) or in the pancreas (PIP, LIP) (**f**) following T/P treatment.

Senescence induction is associated with dynamic transcriptional and chromatin changes. Rb and p53-regulated pathways mediate repression of cell cycle genes^34,35^. In addition, a number of other transcription factors and regulators, including NF-κB, C/EBPβ, cGAS-STING, JAK/STATs, and NOTCH, lead to activation of divergent SASP programs^36–39^. These transcriptional changes are accompanied by dramatic remodeling of the chromatin landscape, with Rb enabling repressive H3K9me3-mediated chromatin compaction at cell cycle genes^35^, and BRD4 facilitating H3K27Ac-mediated enhancer activation at SASP loci^40^. Transcription factor enrichment analysis demonstrated that transcriptional targets of NF-κB and its p65 subunit RELA, which we have shown to be important activators of the SASP following T/P-induced senescence^19,20^, were induced preferentially in tumors propagated in the lung TME (Fig. 2e). Targets of interferon regulatory factors (IRFs) that respond to and drive interferon production downstream of STING pathway activation were also enriched in tumors in the lung as compared to pancreas TME (Fig. 2e). Interestingly, regulators of 3D chromatin topology and DNA looping (CTCF, RAD21, SMC3), as well as histone modifications and chromatin compaction (EZH2, p300), were enriched in tumors within the pancreas TME (Fig. 2f). These findings suggested the possibility that chromatin remodeling within tumors in the pancreas TME may lead to transcriptional repression of specific SASP genes and regulators.

### EZH2 methyltransferase activity leads to pro-inflammatory SASP suppression in PDAC

EZH2 is a member of the polycomb repressor complex 2 (PRC2) with methyltransferase activity that mediates transcriptional gene repression through H3K27 trimethylation (H3K27me3)^41^. Suppression of EZH2 can induce cellular senescence and a subsequent SASP in fibroblasts through activation of DNA-damage response (DDR) pathways and loss of repressive H3K27me3 marks on *CDKN2A* (i.e. p16), a canonical regulator of senescence-associated cell cycle arrest, and other SASP gene loci^42–44^. In addition,

EZH2 induction in other cancer settings mediates suppression of NK and T cell immune surveillance and immunotherapy resistance^45–48^. We found expression of EZH2 repressed genes significantly decreased, and global levels of H3K27me3 dramatically increased, in tumors propagated in the pancreas as compared to those in the lungs (Extended Data Fig. 4a,b). Based on these findings, we hypothesized that EZH2 activation may lead to epigenetic silencing of the SASP through its methyltransferase activity that contributes to suppression of cytotoxic lymphocyte immunity in PDAC.

Short hairpin RNAs (shRNAs) were generated that could potently suppress levels of EZH2 or SUZ12, another PRC2 complex component that interacts with EZH2, in our *KPC* PDAC cell lines (Fig. 3a). EZH2 knockdown had no impact on senescence-induced growth arrest or expression of SA-β-gal or other senescence-related genes (e.g. *Cdkn2a*, *Cdkn2b*) following T/P treatment in *KPC* PDAC cells (Extended Data Fig. 4c,d). In contrast, SASP-related pro-inflammatory cytokines (e.g. IL-6, IL-15, IL-18) and chemokines (CCL2/5/7/8/20, CXCL2/10) important for cytotoxic lymphocyte immunity were significantly upregulated at the gene expression and protein secretion levels in *KPC* PDAC cells harboring *Ezh2* or *Suz12*-targeting shRNAs as compared to those harboring a control *Renilla* (*Ren*) shRNA following T/P treatment (Fig. 3b and Extended Data Fig. 4d). Immunomodulatory cell surface proteins associated with the SASP, including cell adhesion molecules (ICAM-1) important for tumor-lymphocyte synapses, stress ligands that bind and stimulate the activating NKG2D receptor on NK cells (RAETs, ULBP1, H60s), and MHC-I necessary for antigen presentation to T cells, were also strongly upregulated following therapy-induced senescence and EZH2 knockdown (Extended Data Fig. 4d,e).

**Fig. 3.**
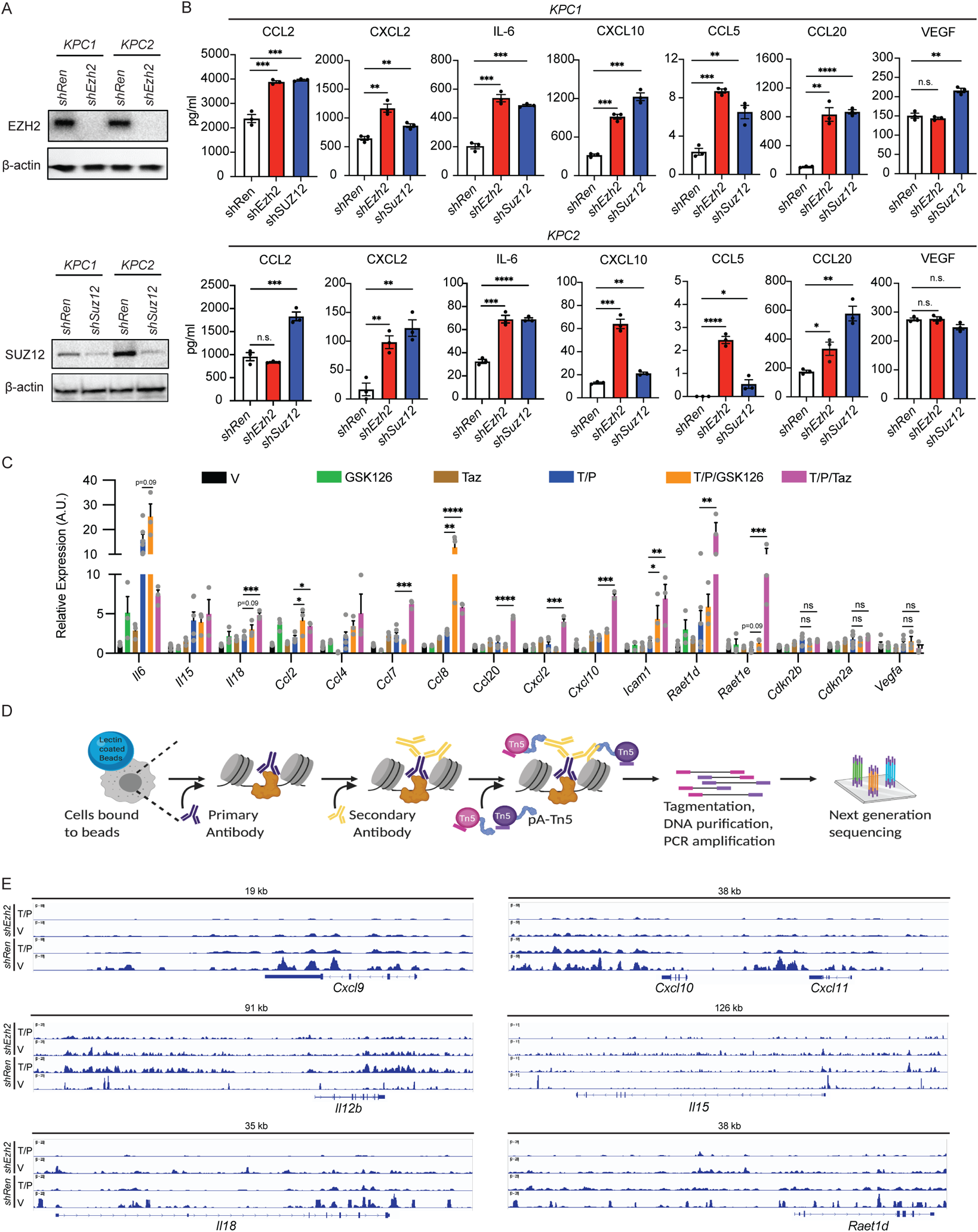
EZH2 methyltransferase activity leads to pro-inflammatory SASP suppression in PDAC. **a**, Immunoblots of *KPC1* and *KPC2* PDAC cells harboring *Renilla* (*Ren*), *Ezh2*, or *Suz12* shRNAs. **b**, Cytokine array results from *KPC1* and *KPC2* PDAC cells with indicated shRNAs treated for 8 days with vehicle or trametinib (25 nM) and palbociclib (500 nM) (n=3). **c**, qRT-PCR analysis of senescence and SASP gene expression in *KPC1* PDAC cells treated with vehicle, trametinib (25 nM), palbociclib (500 nM), GSK126 (1 μM), and/or tazemetostat (5 μM) for 8 days (n=3-6 per group). A.U., arbitrary units. **d**, Schematic of the CUT&Tag protocol used in (**e**). **e**, Genome browser tracks from CUT&Tag analysis showing H3K27me3 occupancy at select pro-inflammatory SASP gene loci in *KPC1* PDAC cells harboring *Ren* or *Ezh2* shRNAs treated with vehicle or trametinib (25 nM) and palbociclib (500 nM) for 8 days (n=2-4 per group). *P* values in **b**-**c** were calculated using two-tailed, unpaired Student’s t-test. Error bars, mean ± SEM. **** *P* <0.0001, *** *P* <0.001, ** *P* <0.01, * *P* <0.05. n.s., not significant.

This increased expression of inflammatory molecules observed following EZH2 inhibition was dependent on EZH2 methyltransferase activity, as treatment with the well-characterized EZH2 methyltransferase inhibitors GSK126 and tazemetostat (Taz) induced pro-inflammatory SASP factors and immunomodulatory cell surface proteins without impacting senescence-associated cell cycle arrest to a similar extent as EZH2 genetic knockdown (Fig. 3c and Extended Data Fig. 4e,f). Pharmacological EZH2 methyltransferase inhibition also led to increased expression of pro-inflammatory SASP and cell surface molecules following T/P-induced senescence in the human PDAC cell line PANC-1 (Extended Data Fig. 4g). Thus, EZH2 blockade leads to enhanced pro-inflammatory SASP production following therapy-induced senescence in PDAC.

To determine whether chromatin compaction governed by histone modifications contributes directly to SASP gene reprogramming following EZH2 targeting, we performed CUT&Tag analysis^49^ to assess the impact of EZH2 suppression on H3K27me3 occupancy at SASP gene loci (Fig. 3d). This analysis uncovered synergistic H3K27me3 loss at the loci of specific pro-inflammatory SASP and NK cell ligand genes, including *Cxcl9*, *Cxcl10, Cxcl11, Il12*, *Il15*, *Il18*, and *Raet1d*, following combined EZH2 knockdown and T/P-induced senescence (Fig. 3e and Supplementary Tables 2-6). In contrast, whereas H3K27me3 marks at pro-angiogenic SASP genes such as *Vegfa, Pdgfa, Pdgfb,* and *Mmp9* were reduced following T/P treatment in the control *shRen* setting, H3K27me3 peaks remained unchanged or even increased at these loci following T/P treatment in the *shEzh2* setting (Extended Data Fig. 4h and Supplementary Tables 2-6), suggesting EZH2 suppression preferentially impacted pro-inflammatory SASP genes. Indeed, VEGF production was not stimulated by EZH2 suppression (Fig. 3b-c and Extended Data Fig. 4d). Therefore, suppression of EZH2 methyltransferase activity in combination with therapy-induced senescence can synergistically reverse the epigenetic repression and promote the transcriptional activation of specific pro-inflammatory SASP genes in PDAC.

### EZH2 blockade activates NK and T cell-mediated long-term tumor control following therapy-induced senescence in PDAC models

To understand the impact of EZH2 suppression on senescence-mediated anti-tumor immunity in PDAC, we transplanted *KPC1* or *KPC2* PDAC cells harboring control Renilla (*shRen*) or EZH2-targeting (*shEzh2*) shRNAs orthotopically into the pancreas of C57BL/6 mice (Fig. 4a). Transplanted *shEzh2* PDAC cells formed tumors at a similar rate as *shRen* PDAC cells and maintained EZH2 knockdown *in vivo* (Extended Data Fig. 5a). Following tumor formation as determined by ultrasound imaging, mice were randomized into treatment groups where they received T/P or a vehicle control (Fig. 4a).

**Fig. 4.**
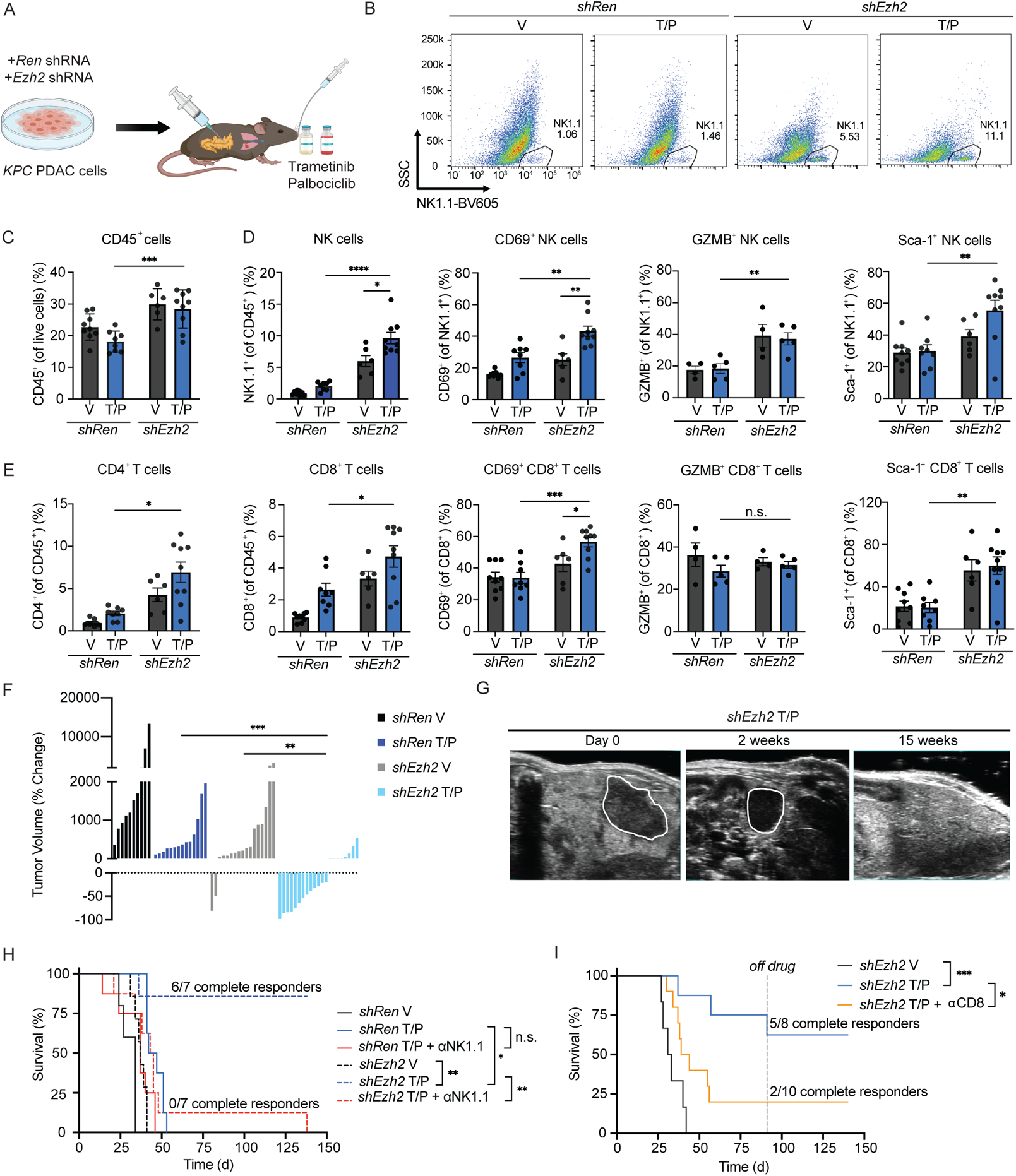
EZH2 blockade activates NK and T cell-mediated long-term tumor control following therapy-induced senescence in PDAC models. **a**, Schematic of *KPC* PDAC syngeneic orthotopic transplant model and treatment regimens. **b**, Representative flow cytometry plots of CD45^+^CD3^−^NK1.1^+^ NK cells in *KPC1* orthotopic PDAC tumors harboring indicated shRNAs from mice treated with vehicle (V) or combined trametinib (1mg/kg) and palbociclib (100mg/kg) (T/P) for 2 weeks. SSC, side scatter. **c-e**, Flow cytometry analysis of total CD45^+^ immune cells (**c**), NK cell numbers and activation markers (**d**), and T cell numbers and activation markers (**e**) in *KPC1* orthotopic PDAC tumors harboring indicated shRNAs following treatment as in (**b**) (n ≥ 4 per group). **f**, Waterfall plot of the response of *KPC1* orthotopic PDAC tumors with indicated shRNAs to treatment as in (**b**) (n ≥ 10 per group). **g**, Representative ultrasound images of *shEzh2 KPC*1 orthotopic PDAC tumors prior to treatment and after 2 or 15 weeks of treatment with combined trametinib (1 mg/kg) and palbociclib (100 mg/kg). PDAC tumors are outlined in white. **h**, Kaplan-Meier survival curve of mice with *KPC*1 orthotopic PDAC tumors harboring indicated shRNAs treated with vehicle, combined trametinib (1mg/kg) and palbociclib (100mg/kg), and/or an NK1.1 depleting antibody (PK136; 250 μg) (n ≥ 5 per group). **i,** Kaplan-Meier survival curve of mice with *shEzh2 KPC*1 orthotopic PDAC tumors treated with vehicle, combined trametinib (1mg/kg) and palbociclib (100mg/kg), and/or a CD8 depleting antibody (2.43; 200 μg) (n ≥ 6 per group). Dotted line indicates timepoint when mice were taken off of treatment. *P* values in **c-f** were calculated using two-tailed, unpaired Student’s t-test, and those in **h** and **i** calculated using log-rank test. Error bars, mean ± SEM. **** *P* <0.0001, *** *P* <0.001, ** *P* <0.01, * *P* <0.05. n.s., not significant.

Immunophenotyping by multi-parametric flow cytometry analysis following two-week treatment revealed significant changes in lymphocyte numbers and activity. T/P treatment in the setting of EZH2 knockdown led to increased total leukocyte infiltration, including enhanced NK and CD4^+^ and CD8^+^ T cell accumulation (Fig. 4b-e and Extended Data Fig. 5b,c). NK cells and CD8^+^ T cells also expressed higher levels of early activation (CD69, Sca-1) markers, and NK cells (but not CD8^+^ T cells) appeared more cytotoxic by expression of GZMB following T/P treatment of *shEzh2* as compared to control *shRen KPC1*-derived tumors (Fig. 4d,e and Extended Data Fig. 5b,c). This increase in cytotoxic lymphocytes following T/P-induced senescence and EZH2 blockade was also accompanied by a decrease in F4/80^+^ macrophages (Extended Data Fig. 5d).

Combinatorial EZH2 knockdown and T/P treatment also had profound anti-tumor effects. Whereas T/P treatment in the control *shRen* setting or EZH2 knockdown alone led to a marginal reduction in tumor growth, T/P treatment in the context of EZH2 knockdown produced significant tumor control, with many tumors regressing after just two-week treatment (Fig. 4f,g and Extended Data Fig. 5e). Remarkably, the majority of *shEzh2 KPC1*-derived tumors treated with T/P continued to regress and completely responded (Fig. 4g-i). Indeed, while mice harboring control *shRen* PDAC treated with vehicle or T/P or *shEzh2* PDAC treated with vehicle quickly succumbed to the disease, 11/15 mice harboring *shEzh2* tumors treated with T/P had complete responses that remained durable even after treatment ceased (Fig. 4h,i). *KPC2* PDAC transplant mice also showed enhanced survival following EZH2 knockdown and treatment with T/P, including 3/8 complete responders (Extended Data Fig. 5f).

We treated some PDAC-bearing mice with NK1.1 (PK136) or CD8 (2.43) depleting monoclonal antibodies (mAbs) simultaneously with drug administration to assess whether activation of NK and/or CD8^+^ T cell immunity was responsible for tumor control. Strikingly, both NK or CD8^+^ T cell depletion mitigated long-term survival and prevented complete tumor responses induced following T/P treatment of animals with EZH2 suppressed *KPC1* and *KPC2* PDAC tumors (Fig. 4h,i and Extended Data Fig. 5f). Together these findings demonstrate that EZH2 knockdown can potentiate senescence-mediated long-term tumor control in PDAC through mobilization of cytotoxic NK and T lymphocyte immunity.

### EZH2 suppression reinstates SASP-associated chemokines to drive NK and T cell accumulation in PDAC

Given the numerous cell autonomous and non-cell autonomous functions of EZH2 in cancer biology^50^, we performed bulk RNA-seq on FACS sorted GFP^+^ tumor cells isolated from drug-treated *shRen* or *shEzh2* PDAC tumors to understand the mechanisms by which EZH2 targeting led to immune-mediated tumor control. Unbiased KEGG pathway analysis revealed “cytokine-cytokine receptor interaction” and “cell adhesion molecules” as top differentially regulated pathways when comparing *shEZH2* vs. *shRen* tumors treated with T/P (Fig. 5a). Deeper analysis revealed significantly increased expression of SASP-associated pro-inflammatory cytokines and chemokines (*Il15, Ccl2/5/7/8, Cxcl9/10/11, Cx3cl1)*, as well as genes involved in antigen presentation/processing (*B2m, Tap1, Tapbp*) and cell adhesion (*Icam1*) important for T and NK cell recognition of tumor cells in the context of T/P-induced senescence and EZH2 suppression (Fig. 5b and Supplementary Table 7).

**Fig. 5.**
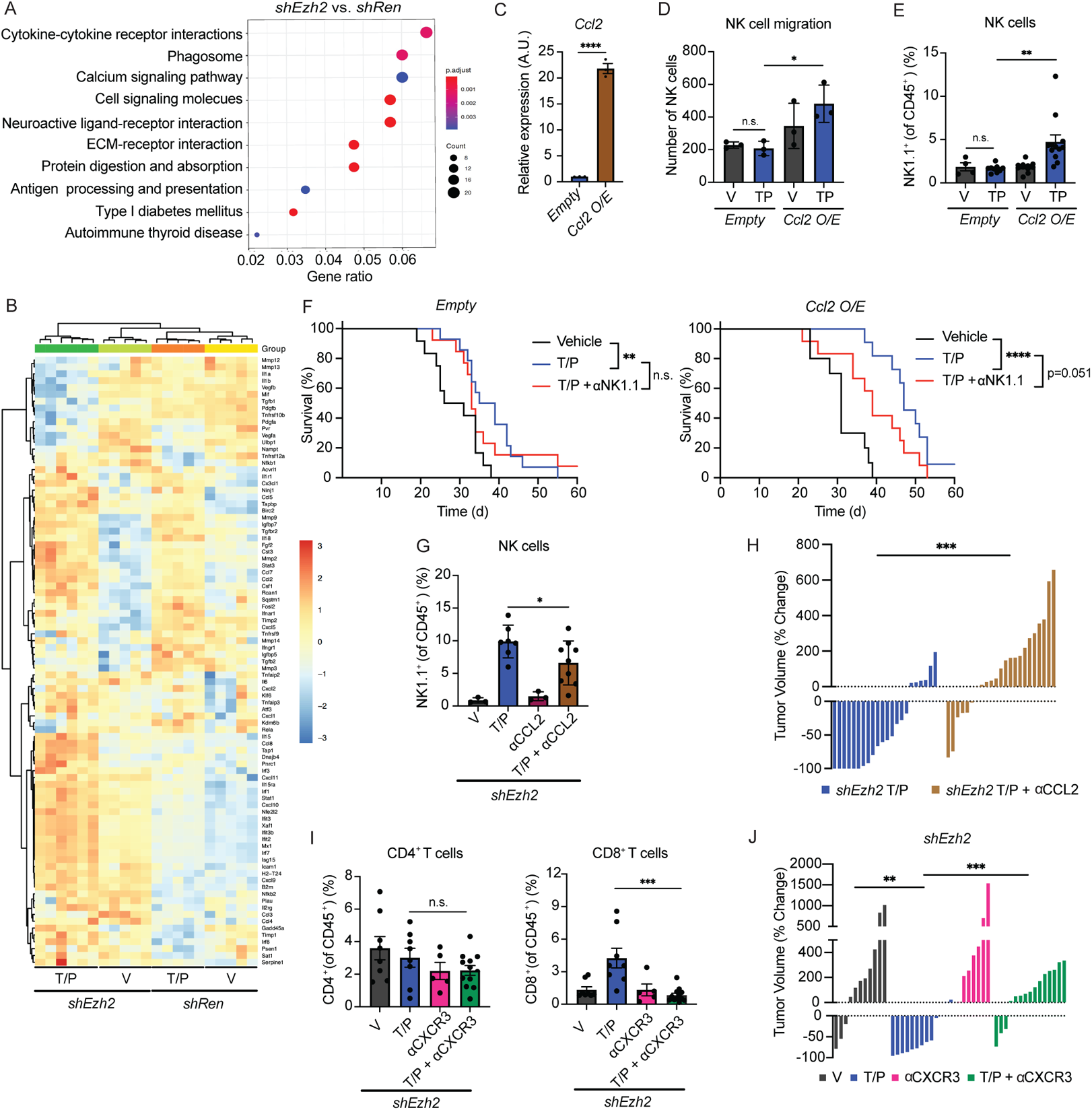
EZH2 suppression reinstates SASP-associated chemokines to drive NK and T cell accumulation in PDAC. **a**, KEGG pathway analysis of RNA-seq data showing top enriched pathways in *shEzh2* as compared to *shRen KPC*1 orthotopic PDAC tumor cells in mice treated with combined trametinib (1mg/kg) and palbociclib (100mg/kg) for 2 weeks (n=5-6 per group). **b**, Heatmap of RNA-seq analysis of SASP gene expression in tumor cells sorted from *KPC*1 orthotopic PDAC tumors harboring indicated shRNAs and treated with vehicle or combined trametinib (1mg/kg) and palbociclib (100mg/kg) for 2 weeks (n=5-6 per group). **c**, qRT-PCR analysis of *Ccl2* expression in *KPC1* PDAC cells engineered to overexpress (O/E) a *Ccl2* cDNA or *Empty* control vector (n=3). A.U., arbitrary units. **d**, NK cell migration assay in the presence of conditioned media from *KPC1* PDAC cells engineered to overexpress *Ccl2* or an *Empty* control vector and treated with vehicle or trametinib (25nM) and palbociclib (500nM) for 8 days (n=3). **e**, Flow cytometry analysis of NK cell numbers in *KPC*1 orthotopic PDAC tumors expressing control *Empty* or *Ccl2* vectors following treatment as in (**b**) (n ≥ 4 per group). **f**, Kaplan-Meier survival curve of mice with *KPC*1 orthotopic PDAC tumors expressing control *Empty* (left) or *Ccl2* (right) vectors treated with vehicle, combined trametinib (1mg/kg) and palbociclib (100mg/kg), and/or an NK1.1 depleting antibody (PK136; 250 μg) (n ≥ 10 per group). **g**, Flow cytometry analysis of NK cell numbers in *shEzh2 KPC*1 orthotopic PDAC tumors following treatment with vehicle, combined trametinib (1mg/kg) and palbociclib (100mg/kg), and/or a CCL2 depleting antibody (2H5; 200 μg) for 2 weeks (n ≥ 3 per group). **h**, Waterfall plot of the response of *shEzh2 KPC1* orthotopic PDAC tumors to treatment as in (**g**) (n ≥ 22). **i**, Flow cytometry analysis of CD4^+^ and CD8^+^ T cell numbers in *shEzh2 KPC*1 orthotopic PDAC tumors following treatment with vehicle, combined trametinib (1mg/kg) and palbociclib (100mg/kg), and/or a CXCR3 depleting antibody (CXCR3-173; 200 μg) for 2 weeks (n ≥ 5 per group). **j**, Waterfall plot of the response of *shEzh2 KPC1* orthotopic PDAC tumors to treatment as in (**i**) (n ≥ 7 per group). *P* values in **c-e** and **g-j** were calculated using two-tailed, unpaired Student’s t-test, and those in **f** calculated using log-rank test. Error bars, mean ± SEM. **** *P* <0.0001, *** *P* <0.001, ** *P* <0.01, * *P* <0.05. n.s., not significant.

The expression of other SASP-associated factors showed differential responses to EZH2 blockade. Whereas transcriptional regulators of the pro-inflammatory SASP that are normally repressed in the PDAC TME, including STING (*Irf1/3/7/8, Ifnar, Isg15*) and STAT (*Stat1/3*) pathway components, were upregulated following therapy-induced senescence and EZH2 suppression (Fig. 5b), many of the pro-angiogenic (*Vegfa/b, Pdgfa/b, Mmp3/9/12/13/14*) and immune suppressive (*Tgfb1/2, Cxcl1/5*) SASP factors normally induced during senescence were downregulated (Fig. 5b). Indeed, the increase in blood vessels observed following T/P treatment in *shRen* tumors and as reported in our previous study^20^ was not found in s*hEzh2* tumors (Extended Data Fig. 6a). Thus, EZH2 suppression following T/P-induced senescence triggers a phenotypic switch in the SASP program in the PDAC TME from a pro-angiogenic SASP to a pro-inflammatory SASP that may contribute to enhanced cytotoxic lymphocyte anti-tumor immunity.

Many of the most highly induced pro-inflammatory SASP factors in the lung TME (Fig. 2c,d) or upon EZH2 knockdown in the PDAC TME (Fig. 5b) following therapy-induced senescence are chemokines known to attract NK and T cells from the periphery into inflamed tissues^33^, including CCL2 and CXCL9/10. In addition to its impact on monocyte and macrophage trafficking, CCL2 can also attract NK cells expressing its receptor CCR2^51^, as has been previously shown in senescent liver cancer lesions^52^. To interrogate the role of CCL2 in anti-tumor NK cell responses in PDAC, we first engineered *KPC* PDAC cell lines to express a *Ccl2* cDNA (or an *Empty* vector as a control) (Fig. 5c). *In vitro*, conditioned media from *KPC1* cells overexpressing CCL2 and pre-treated with T/P produced significantly more NK cell migration through a transwell insert (Fig. 5d), demonstrating that CCL2 secretion by senescent tumor cells can facilitate NK cell chemotaxis. *Empty* or *Ccl2* expressing *KPC* cells were then transplanted orthotopically into the pancreas of C57BL/6 mice and mice randomized into treatment groups following tumor formation to assess the impact on NK cell immune surveillance *in vivo*. Flow cytometry analysis revealed that tumor-specific CCL2 overexpression in the context of T/P-induced senescence was sufficient to significantly increase NK cell accumulation into PDAC without affecting NK cell activation (Fig. 5e and Extended Data Fig. 6b). This increased NK cell influx into CCL2 overexpressing tumors prolonged the survival of PDAC-bearing animals treated with T/P, as NK cell depletion with an NK1.1-targeting mAb significantly diminished the survival advantage (Fig. 5f).

To determine whether EZH2 suppression facilitated anti-tumor NK cell immunity through CCL2 induction, we also treated mice transplanted with *shEzh2 KPC1* PDAC tumors with vehicle, T/P, and/ or a mAb targeting CCL2 (2H5). Indeed, CCL2 was required for these effects, as CCL2 blockade resulted in reduced NK cell accumulation and abolished tumor regressions (Fig. 5g,h). Thus, SASP-associated CCL2 is both necessary and sufficient to drive NK cell infiltration and potentiate NK cell-mediated tumor control in PDAC following T/P-induced senescence.

In contrast to its effects on NK cells, CCL2 overexpression or neutralization had little impact on CD4^+^ and CD8^+^ T cell recruitment (Extended Data Fig. 6c,d), suggesting other SASP chemokines may affect T cell chemotaxis. T cells express the receptor CXCR3 that binds chemokines CXCL9/10/11 that are critical for CD8^+^ T cell homing to the TME and ICB immunotherapy efficacy^53–56^. In agreement, treatment of *shEzh2* PDAC-bearing mice with vehicle, T/P, and/or a CXCR3 mAb (CXCR3-173) revealed that CXCR3 blockade blunted CD8^+^ and to a lesser extent CD4^+^ T cell accumulation without affecting NK cell numbers (Fig. 5i and Extended Data Fig. 6e). This reduction in CD8^+^ T cell recruitment upon CXCR3 blockade also mitigated the anti-tumor effects of combined EZH2 suppression and T/P-induced senescence and reversed PDAC tumor regressions (Fig. 5j). Therefore, distinct SASP chemokines are important for NK and T cell chemotaxis and migration into the PDAC TME and enable the immune-mediated anti-tumor effects observed upon EZH2 suppression.

### Pharmacological EZH2 methyltransferase inhibition in combination with T/P reactivates cytotoxic NK and T cell immunity and enhances tumor control in preclinical PDAC models

Given the profound anti-tumor effects of genetic EZH2 knockdown in combination with therapy-induced senescence in PDAC, we next tested whether small molecule EZH2 inhibitors that are in clinical development could achieve similar effects. Mice transplanted orthotopically with *KPC1* PDAC cells were randomized and treated with vehicle, T/P, and/or the FDA-approved EZH2 methyltransferase inhibitor tazemetostat (Taz) at doses reported to potently inhibit H3K27me3 levels *in vivo* (Fig. 6a,b). Short-term two-week treatment with Taz in combination with T/P significantly reduced tumor growth compared to each treatment arm alone (Fig. 6c), though tumor regressions were not as frequent or robust as those found with genetic EZH2 knockdown (Fig. 4f). Combined Taz and T/P treatment also led to a significant increase in NK cell numbers and their expression of activation (e.g. Sca-1) and cytotoxicity (e.g. GZMB) markers (Fig. 6d). Interestingly, whereas CD4^+^ and CD8^+^ T cell numbers increased at lower doses of Taz (125 mg/kg) in combination with T/P, high Taz concentrations (400 mg/kg) reduced CD8^+^ T cell numbers and expression of CD69, a marker of early activation and proliferation (Fig. 6d,e). This suggests that, separate from its action on tumor cells, Taz may affect T cell proliferation in a manner that reduces its anti-tumor activity. We therefore performed the remaining studies using the lower 125 mg/kg Taz dose.

**Fig. 6.**
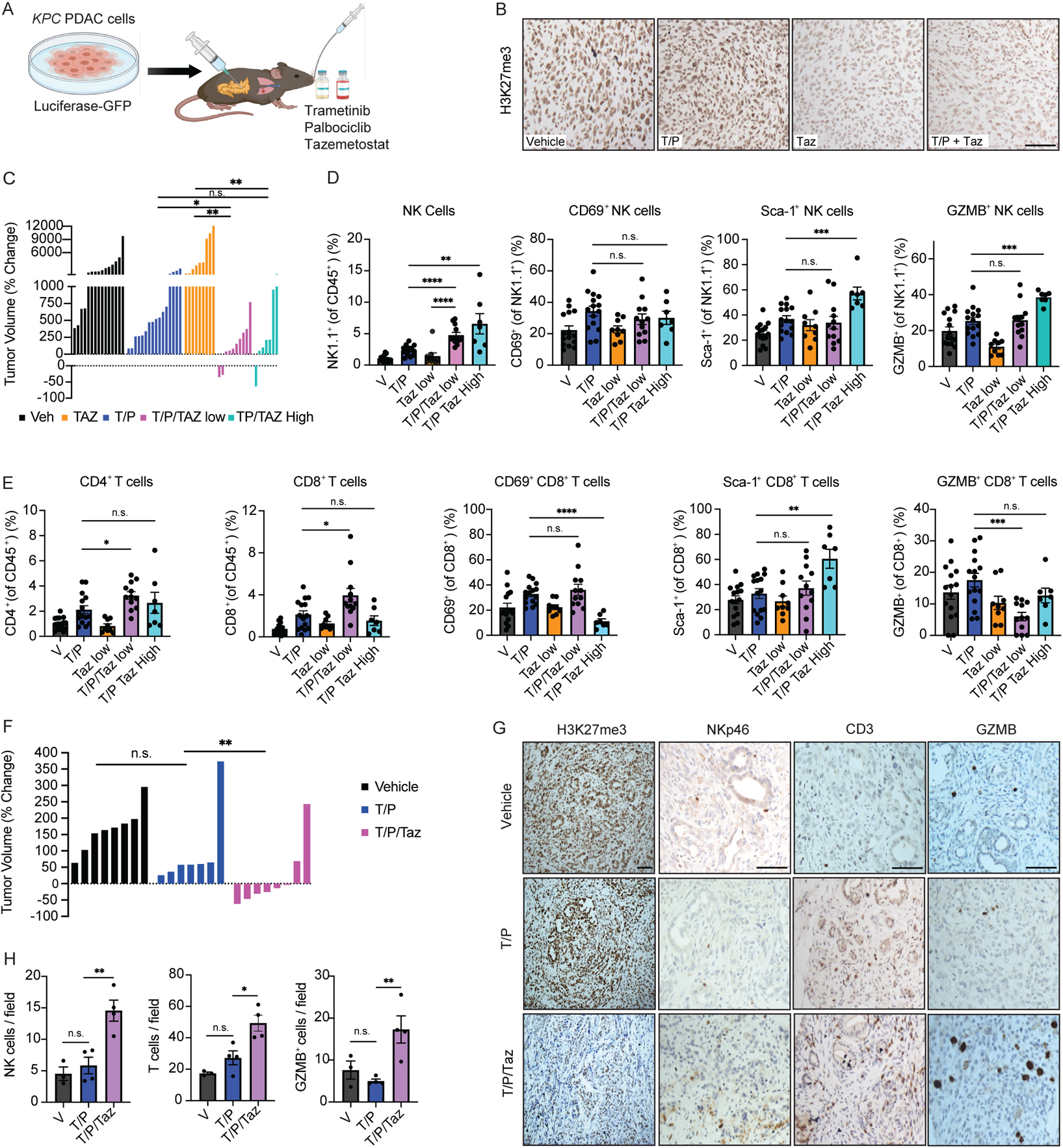
Pharmacological EZH2 methyltransferase inhibition in combination with T/P reactivates cytotoxic NK and T cell immunity and enhances tumor control in preclinical PDAC models. **a**, Schematic of *KPC* PDAC syngeneic orthotopic transplant model and treatment regimens. **b**, Immunohistochemical (IHC) staining of *KPC1* orthotopic PDAC tumors treated vehicle, trametinib (1 mg/kg) and palbociclib (100 mg/kg), and/or tazemetostat (Taz) (125 mg/kg) for 2 weeks. Scale bars, 50μm. **c**, Waterfall plot of the response of *KPC1* orthotopic PDAC tumors following 2 week-treatment with vehicle, trametinib (1 mg/kg) and palbociclib (100 mg/kg), and/or low (125 mg/kg) or high (400 mg/kg) doses of tazemetostat (n ≥ 7 per group). **d-e**, Flow cytometry analysis of NK (**d**) and T cell (**e**) numbers and activation markers in *KPC1* orthotopic PDAC tumors following treatment as in (**c**) (n ≥ 7 per group). **f**, Waterfall plot of the response of *KPC* GEMM tumors to treatment as in (**b**) (n ≥ 7 per group). **g**, IHC staining of *KPC* GEMM tumors treated as in (**b**). Scale bars, 50μm. **h**, Quantification of NKp46^+^ NK cells, CD3^+^ T cells, and GZMB^+^ cells in (**g**) (n ≥ 3 per group). *P* values in **c-f** and **h** were calculated using two-tailed, unpaired Student’s t-test. Error bars, mean ± SEM. **** *P* <0.0001, *** *P* <0.001, ** *P* <0.01, * *P* <0.05. n.s., not significant.

To test this inhibitor combination in an autochthonous model, we utilized *P48-Cre*; *LSL-KRAS^G12D^*;*Trp53^fl/wt^*(*KPC*) GEMM mice that spontaneously develop PDAC that closely resembles the human disease^57^. Two-week combined Taz and T/P treatment of tumor-bearing *KPC* GEMMs led to significantly reduced H3K27me3 levels and tumor regressions in 4/8 mice, in contrast to T/P treatment alone where tumor regressions were not observed (Fig. 6f,g). In addition, Taz treatment in combination with T/P promoted the infiltration of NK cells and enhanced cytotoxic GZMB expression, as well as further increased the T cell accumulation observed with single arm T/P regimens (Fig. 6g,h). Thus, both genetic EZH2 suppression and EZH2 methyltransferase inhibitor treatment can augment T/P-induced senescence to potentiate cytotoxic NK and T cell immunity and tumor control in transplanted and GEMM PDAC models.

### EZH2 is associated with suppression of inflammatory chemokine signaling, reduced NK and T cell immune surveillance, and poor survival in PDAC patients

Finally, we set out to evaluate the relationship between EZH2 activity, inflammatory signaling, and NK and T cell immunity in human PDAC. We first interrogated a previously published gene expression dataset containing PDAC patient samples^58^. EZH2 and PRC2 repression signatures correlated positively with inflammatory response genes, including expression of *CCL2, CXCL9*, and *CXCL10* that are important of NK and T cell trafficking into the PDAC TME (Fig. 7a and Supplementary Table 8). Moreover, NK and CD8^+^ T cell gene transcript levels were also significantly associated with EZH2 repressed gene expression (Fig. 7a and Supplementary Table 8). These results support a relationship between EZH2 activity, inflammatory chemokine signaling, and cytotoxic lymphocyte infiltration in human PDAC.

**Fig. 7.**
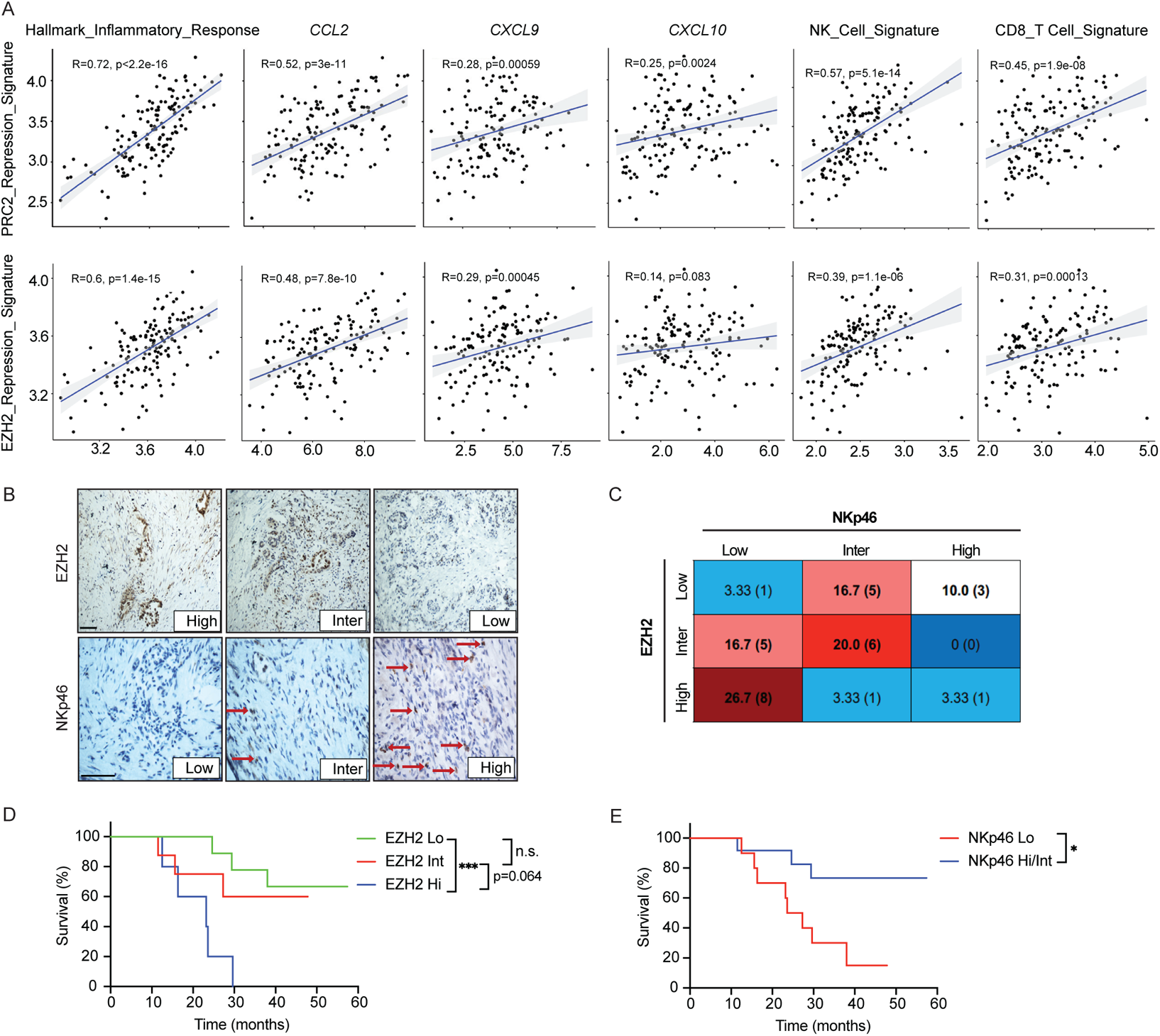
EZH2 is associated with suppression of inflammatory chemokine signaling, reduced NK and T cell immune surveillance, and poor survival in PDAC patients. **a**, Pearson’s correlation analysis plots comparing EZH2 and PRC2 repression signatures with inflammatory response gene sets, *CCL2*, *CXCL9*, and *CXCL10* expression, and NK and CD8^+^ T cell signatures in human primary PDAC transcriptomic data^58^ (n=145 samples). **b**, Representative IHC staining of surgically resected human PDAC tumors (n=30). Arrows indicate NK cells. Scale bars, 50μm. **c**, Scoring of EZH2 and NKp46 expression from IHC staining in (b) (n=30). Percentage of samples with indicated EZH2 and NKp46 scores are shown, with the total number of sample in parentheses. **d**, Kaplan-Meier survival curve of human PDAC patients stratified based on EZH2 expression levels in (**b**) (n=9, 8, and 5 for EZH2 Lo, Int, and Hi, respectively). **e**, Kaplan-Meier survival curve of human PDAC patients stratified based on NKp46 expression levels in (**b**) (n=10 and 12 for NKp46 Lo and Hi/Int, respectively). *P* values in **a** were calculated using two-tailed, unpaired Student’s t-test, and those in **d** and **e** were calculated using log-rank test. Error bars, mean ± SEM. **** *P* <0.0001, *** *P* <0.001, ** *P* <0.01, * *P* <0.05. n.s., not significant.

We then performed immunohistochemical (IHC) staining on formalin-fixed, paraffin-embedded (FFPE) surgically resected tumor specimens from PDAC patients treated at UMass Memorial hospital to assess expression of EZH2 and the NK cell marker NKp46. The 30 patient samples analyzed presented a spectrum of EZH2 expression and NK cell density, with a trend toward low NK cell numbers correlating with high/intermediate EZH2 expression and higher NK cell penetrance correlating with low EZH2 expression (Fig. 7b,c). Patients with low EZH2 expression in their primary PDAC lesions also had significantly increased overall survival compared to patients with high EZH2 PDAC expression (Fig. 7d). Similarly, overall survival was significantly improved in patients with high/intermediate NK cell numbers compared to those with low or absent tumor NK cells (Fig. 7e). Taken together, our work demonstrates that EZH2 is associated with suppression of inflammatory signaling, NK and T cell dysfunction, and reduced survival in murine and human PDAC, and that targeting EZH2 activity can restore long-term innate and adaptive immune-mediated PDAC control.

## DISCUSSION

PDAC remains without durable chemo-, targeted, and immunotherapy regimens, and as such has a dismal 5-year survival rate of 11%^1^. Many promising studies and clinical trials have focused on overcoming immune suppression as a therapeutic strategy in PDAC through (a) re-engineering T cell responses via CAR-T, ICB therapy, or neo-antigen vaccine approaches, (b) targeting suppressive fibroblast populations and functions, and/or (c) eliminating or reprogramming suppressive myeloid cells^59^. Here we investigated how to remodel the tumor secretome directly as a strategy to enhance tumor immunogenicity and transform the immune suppressive PDAC TME. By comparing the effects of therapy-induced senescence on KRAS mutant LUAD and PDAC tumors, we uncovered an epigenetic mechanism that suppresses the pro-inflammatory SASP secretome in PDAC that is mediated by PRC2 component EZH2 and its methyltransferase activity. EZH2 inhibition in combination with therapy-induced senescence unleashes pro-inflammatory SASP chemokines such as CCL2 and CXCL9/10 and induces MHC-I and NK ligand expression to orchestrate an innate and adaptive immune attack through cytotoxic NK and T lymphocytes that in some cases led to complete responses in preclinical PDAC models.

EZH2 has been shown to facilitate tumor immune evasion and resistance to ICB therapy in other cancer settings^45–48^. Here we uncovered a novel mechanism by which EZH2 mediates PDAC immune suppression through inhibition of the pro-inflammatory transcriptome, secretome, and surfaceome associated with the SASP. Mechanistically, EZH2 methyltransferase activity was directly responsible for suppressing many SASP components, such that genetic or pharmacological inhibition of EZH2 in tumor cells triggered to senescence following T/P therapy led to a marked reduction in H3K37me3 marks and increased transcription of many SASP cytokines and chemokines. Clinically, EZH2 is commonly overexpressed in poorly differentiated PDAC and associated with chemoresistance^60,61^. Our analysis of patient samples in addition reveals that EZH2 is not only associated with suppression of inflammatory chemokines and NK and T cell immunity in the human disease, but also poor overall patient survival. Thus, our work reveals EZH2 as an important marker and inducer of immune suppression in PDAC that is therapeutically targetable.

The SASP is often considered a “double-edged sword” and can promote anti-tumor immune surveillance or alternatively pro-tumor immune evasion depending on the context^21,22,28^. Here we find that the resident tissue or TME context plays a key role in immune responses to senescence stimuli. Unexpectedly, we observed a distinct SASP program induced in the pancreas compared to lung TME following T/P-induced senescence that contributed to a lack of NK cell immunity in PDAC. Whereas LUAD or PDAC tumors propagated in the lungs displayed induction of pro-inflammatory SASP factors (e.g. IL-6, IL-15, CCL2, CXCL9/10), tumors in the pancreas expressed high levels of pro-angiogenic SASP factors (e.g. VEGFs, PDGFs, MMPs). Increased EZH2 activity and H3K27me3 levels appeared to mediate this phenotypic switch, as EZH2 blockade led to induction of pro-inflammatory SASP while simultaneously reducing angiogenic SASP factor expression following T/P treatment. Though our current study did not address how the pancreas TME per se inhibits the SASP or upregulates EZH2 activity in tumor cells, we postulate that stromal populations that are prominent in the pancreas TME play a crucial role. Nonetheless, this study demonstrates that the quantity and quality of the SASP elicited following senescence induction is influenced in part by the TME and its impact on the epigenetic state of the cancer cell.

Our findings suggest that induction of chemokines through the SASP that drive NK and T cell trafficking into TMEs could be a powerful approach to make immunologically “cold” tumors such as PDAC “hot”^62^. While often considered strictly a monocyte chemoattract, our results support the emerging view that CCL2 is also an important stimulator of NK cell chemotaxis into senescent tumors^52^. Chemokines CCL7 and CCL8 are also highly induced in senescent PDAC cells following EZH2 inhibition and bind to the same receptor as CCL2 (CCR2), suggesting they could also play a role in NK cell trafficking into PDAC. Other SASP chemokines such as CXCL9/10 that bind to CXCR3 and are associated with T cell recruitment and “hot” TMEs in other cancer settings^53,54,56,63^ are necessary for CD8^+^ T cell recruitment into PDAC following therapy-induced senescence. Remarkably, in our system, combining increased cytotoxic NK and CD8^+^ T cell trafficking via SASP chemokines with the enhanced immunogenicity of senescent cells is sufficient to potentiate anti-tumor immune surveillance in PDAC even in the absence of immune checkpoint blockade.

Though both genetic EZH2 suppression and pharmacological EZH2 methyltransferase inhibition lead to NK and T cell activation and enhanced PDAC tumor control following therapy-induced senescence, the anti-tumor effects of small molecule EZH2 inhibition are not as robust or durable as compared with its genetic knockdown. The scaffolding functions of EZH2, acting independently of its methyltransferase activity, may contribute to the more potent effects of EZH2 protein suppression^64,65^. Inhibitors that disrupt PRC2 complex stability, for instance by targeting the core subunit EED^66,67^, or EZH2 degraders^68–70^, may offer more potent PRC2 complex and EZH2 suppression. Alternatively, systemic inhibition of EZH2 using small molecule inhibitors undoubtedly impacts non-tumor cells as well, and their deleterious effects on T cell function and proliferation could reduce their anti-tumor activity^71^. Here, further optimization of dose and scheduling may help improve the therapeutic index of EZH2 targeted therapies.

Tazemetostat and other EZH2 methyltransferase inhibitors have demonstrated efficacy and been implemented into the clinical care of hematological malignancies and sarcomas; however, they have yet to show potent activity as single agents in solid tumors^72^. Our findings provide rationale for combining EZH2 inhibitors with a senescence-inducing therapy – here produced by a MEK and CDK4/6 inhibitor combination – to promote NK and T cell-mediated eradication of senescent PDAC lesions through pro-inflammatory SASP induction. As radiation and chemotherapy can also induce senescence in some settings, it will be interesting to see whether EZH2 inhibitors show combinatorial activity with these agents as well. Collectively, our work provides a novel strategy for leveraging EZH2 inhibitors as an immune oncology approach in combination with senescence-inducing agents to remodel the inflammatory tumor secretome for immune-mediated PDAC control.

## METHODS

The research performed in this study complies with all ethical regulations. All mouse experiments were approved by the University of Massachusetts Chan Medical School Internal Animal Care and Use Committee (IACUC). Surgically resected PDAC patient samples were acquired under the University of Massachusetts Chan Medical School IRB protocol no. H-4721.

### Cell lines and compounds

PANC-1 cells were purchased from the American Type Culture Collection (ATCC). Murine PDAC (*KPC1, KPC2*) and LUAD (*KP1, KP2*) cell lines were generated as previously described^19,20^. For visualizing and tracking *KPC* and *KP* tumor cell lines with luciferase and GFP *in vivo,* cells were transduced with the following retroviral constructs: MSCV-luciferase (luc)-IRES-GFP (for *KPC1* and *KP1*), MSCV-IRES-GFP (for *KP2*), and MSCV-shRen-PGK-Puro-IRES-GFP (for *KPC2*). Retroviruses were packaged by co-transfection of Gag-Pol expressing 293 T cells with expression constructs and envelope vectors (VSV-G). Following transduction, cells were purified by FACS sorting the GFP^+^ population on a FACSAria (BD Biosciences). All cells were maintained in a humidified incubator at 37°C with 5% CO_2_, and grown in DMEM supplemented with 10% FBS and 100 IU/ml penicillin/streptomycin (P/S). *KPC* cell lines were grown in culture dishes coated with 100 µg/ml collagen (PureCol) (5005; Advanced Biomatrix). All cell lines used were negative for mycoplasma. Human cell lines were authenticated by their source repository.

Trametinib (S2673), palbociclib (S1116), and GSK126 (S7061) were purchased from Selleck chemicals for *in vitro* studies. Drugs for *in vitro* studies were dissolved in DMSO (vehicle) to yield 10mM stock solutions and stored at −80°C. For *in vitro* studies, growth media with or without drugs was changed every 2-3 days. For *in vivo* studies, trametinib (T-8123) and palbociclib (P-7744) were purchased from LC Laboratories, and tazemetostat (HY-13803) purchased from MedChemExpress. Trametinib was dissolved in a 0.5% hydroxypropyl methylcellulose and 0.2% Tween-80 solution, palbociclib in 50 mM sodium lactate buffer (pH 4), and tazemetostat in a 0.5% sodium carboxymethylcellulose and 0.1% Tween-80 solution (Sigma-Aldrich).

### Short-hairpin RNA (shRNA) knockdown

shRNAs targeting *Ezh2*, *Suz12*, and *Renilla (Ren)* were cloned into the XhoI EcoRI locus of MLP retroviral vectors (MSCV-LTR-shRNA-PGK-Puro-IRES-GFP) as previously described^73^. Retroviruses were packaged by co-transfection of Gag-Pol expressing 293 T cells with expression constructs and envelope vectors (VSV-G) using polyethylenimine (PEI; Sigma-Aldrich). Following transduction with shRNA retroviral constructs, cell selection was performed with 4μg/ml puromycin for 1 week. Knockdown was confirmed by Western blot, qRT-PCR, and immunohistochemistry following transplantion into C57BL/6 mice.

### CCL2 overexpression

Murine *Ccl2* cDNA was cloned into an MSCV-based retroviral vector (MSCV-blast). Retroviruses were packaged by co-transfection of Gag-Pol expressing 293 T cells with expression constructs and envelope vectors (VSV-G) using polyethylenimine (PEI; Sigma-Aldrich). Following transduction with *Ccl2* or control *Empty* constructs, cell selection was performed with 10μg/ml Blasticidin S for 1 week. *Ccl2* expression was confirmed by qRT-PCR.

### SA-β-gal staining

SA-β-gal staining was performed as previously described at pH 5.5 for mouse cells and tissue^19,20^. Fresh frozen sections of tumor tissue, or adherent cells plated in 6-well plates, were fixed with 0.5% glutaraldehyde in PBS for 15 min, washed with PBS supplemented with 1mM MgCl_2_, and stained for 4–18 hours in PBS containing 1 mM MgCl_2_, 1mg/ml X-Gal, and 5 mM each of potassium ferricyanide and potassium ferrocyanide. Tissue sections were counterstained with eosin. 5-10 high power 20x fields per tissue section were counted and averaged.

### Drug withdrawal clonogenic assays

*KPC* tumor cells were initially plated in 6-well plates and pre-treated with vehicle (DMSO), trametinib (25 nM), palbociclib (500 nM), and/or tazemetostat (5 μM) for 8 days. Pre-treated cells were then trypsinized, and 5×10^3^ cells re-plated per well of a 6-well plate in the absence of drugs for 7 days. Remaining cells were fixed with methanol (1%) and formaldehyde (1%), stained with 0.5% crystal violet, and photographed using a digital scanner.

### Immunoblotting

Cell lysis was performed using RIPA buffer (Cell Signaling) supplemented with phosphatase inhibitors (5mM sodium fluoride, 1 mM sodium orthovanadate, 1 mM sodium pyrophosphate, 1 mM β-glycerophosphate) and protease inhibitors (Protease Inhibitor Cocktail Tablets, Roche). Protein concentration was determined using a Bradford Protein Assay kit (Biorad). Proteins were separated by SDS-PAGE and transferred to polyvinyl difluoride (PVDF) membranes (Millipore) according to standard protocols. Membranes were immunoblotted with antibodies against EZH2 (5246) and SUZ12 (3737) from Cell Signaling in 5% milk in TBS blocking buffer. After primary antibody incubation, membranes were probed with an ECL anti-rabbit IgG secondary antibody (1:10,000) from GE Healthcare Life Science and imaged using a ChemiDoc imaging system (BioRad). Protein loading was measured using a monoclonal β-actin antibody directly conjugated to horseradish peroxidase (A3854, Sigma-Aldrich) and imaged as above.

### qRT-PCR

Total RNA was isolated using the RNeasy Mini Kit (Qiagen), and complementary DNA (cDNA) was obtained using the TaqMan reverse transcription reagents (Applied Biosystems). Real-time PCR was performed in triplicate using SYBR Green PCR Master Mix (Applied Biosystems) on the StepOnePlus Real-Time PCR system (Applied Biosystems). β-actin or Gapdh served as endogenous normalization controls. qRT-PCR primer sequences can be found in Supplementary Table 9.

### CUT&Tag analysis

CUT&Tag was performed largely as previously described^49^. Trypsinized cells were counted, and 100,000 cells were washed and resuspended in wash buffer (20 mM HEPES pH 7.5; 150 mM NaCl; 0.5 mM Spermidine; 1x Protease inhibitor cocktail) and used for CUT&Tag. 10 µl of activated Concanavalin A coated magnetic beads (Polysciences) were added per sample and incubated at room temperature (RT) for 15 min. Bead-bound cells were resuspended in 100 µl Dig-wash Buffer (20 mM HEPES pH 7.5; 150 mM NaCl; 0.5 mM Spermidine; 0.05% Digitonin; 1x Protease inhibitor cocktail) containing 2 mM EDTA and 1 µl of H3K27me3 antibody (ThermoFisher, MA5-11198). The mixture was incubated overnight at 4 °C for antibodies to bind. After pulling beads to the side of the tube using a magnetic rack and removal of unbound primary antibody, beads were resuspended in 100 µl Dig-wash Buffer containing 1 µl of Guinea Pig anti-Rabbit antibody (Antibodies-Online, ABIN101961) and incubated for 30 mins at RT. Cells were washed 3 times with Dig-wash and then incubated with a 1:50 dilution of pA-Tn5 adapter complex in Dig-med (0.05% Digitonin, 20 mM HEPES, pH 7.5, 300 mM NaCl, 0.5 mM Spermidine, 1x Protease inhibitor cocktail) at RT for 1 hr. Cells were washed thrice in Dig-med Buffer and then resuspended in 300 µl Dig-med Buffer containing 10 mM MgCl_2_ and incubated at 37 °C for 1 hr to activate tagmentation. To stop tagmentation, 10 µl of 0.5 M EDTA, 3 µl of 10% SDS and 1 µl of 20 mg/ml Proteinase K was added to each tube, which were incubated at 55 °C for 1 hr. DNA was extracted by performing one phenol:chloroform extraction followed by ethanol precipitation. The DNA pellet was resuspended in 22 µl of 10 mM Tris pH 8.

CUT&Tag libraries were amplified by mixing 21 µl of tagmented DNA with 2µl each of (10 µM) barcoded i5 and i7 primers^74^, using a different combination for each sample. 25 µl NEBNext HiFi 2x PCR Master mix (NEB) was added to each, and PCR was performed using the following cycling conditions: 72 °C for 5 min (gap filling); 98 °C for 30 s; 17 cycles of 98 °C for 10 s and 63 °C for 30 s; final extension at 72 °C for 1 min and holding at 4 °C. 1.1x volumes of Ampure XP beads (Beckman Coulter) were incubated with libraries for 10 min at RT to clean up the PCR reaction. Bead bound DNA was purified by washing twice with 80% ethanol and eluting in 20 µl 10 mM Tris pH 8.0.

The libraries were quantified by Qubit and paired-end sequencing was performed on an Illumina NextSeq 500 (38 bases for reads 1 and 2 and 8 base indexing on both ends). Paired-end reads were aligned to the mouse reference genome GRCm38 (Ensembl, version 101) using bwa mem^75^ after quality assurance with FastQC (https://www.bioinformatics.babraham.ac.uk/projects/fastqc/). Alignment files in the SAM format were first sorted by coordinates and converted into the BAM format using SAMtools^76^. Subsequently, PCR duplicates were removed from the BAM files using “MarkDuplicates” command of the Picard tools (https://broadinstitute.github.io/picard/). The resulting BAM files were name sorted using SAMtools again. Peaks per condition were called using Genrich (https://github.com/jsh58/Genrich) with name-sorted, de-duplicated BAM files of all biological replicates for a given condition as input and a q-value cutoff of 0.05. Given that H3K27me3 modification are widespread across inactive gene regions, peaks with sizes less than 1 kb were filtered out. Consensus peaks-by-sample count matrix were determined using DiffBind^77^. Differential peak analysis was conducted using DEseq2^78^ with hidden variations adjusted for using svaseq^79^. Peaks with absolute values of log_2_ (shrunken fold change) greater than one and *p*-values less than 0.05, which were corrected for multiple testing using the Benjamini-Hochberg procedure^80^, were considered as significantly differential peaks. Over-representation analysis of differential peaks against custom gene sets and each collection of the MSigDB gene sets^81,82^ was performed using msigdbr (https://github.com/igordot/msigdbr) and clusterProfiler^83^. Track views were generated using the Integrative Genomics Viewer (IGV)^84^.

### Cytokine array

Cells were plated in duplicate or triplicate in 6-well plates and drug treated for 6 days. On day 6, 2 ml of new drug-containing media was added to each well and cells were incubated an additional 48 hours. Conditioned media was then collected and cells trypsinized and counted using a Countess II cell counter (Invitrogen). Media samples were then normalized based on cell number by diluting with culture media. Aliquots (75 μl) of the conditioned media were analyzed using a multiplex immunoassay (Mouse Cytokine/Chemokine 44-Plex array) from Eve Technologies.

### NK cell migration assay

Primary NK cells were isolated and enriched the day of the experiment from the spleens of 8-12 week old female C57BL/6 mice using the NK Cell Isolation Kit II according to manufacturer’s instructions (Miltenyi Biotec). 50,000 NK cells were then seeded in the top chamber of a transwell insert (Corning) in a 24-well dish in serum-Free DMEM media with 100 IU/ml penicillin/streptomycin. Serum-free conditioned media from *KPC* tumor cells (collected for 48 hrs following 6 day pre-treatment with indicated drugs) was then placed in the bottom chamber. Following 4 hr incubation in a 37°C cell culture incubator, NK cells migrating through the bottom chamber were fixed with 4% paraformaldehyde (PFA), stained with DAPI, and counted on a Celigo imaging cytometer (Nexcelom).

### Animal models

All mouse experiments were approved by the University of Massachusetts Chan Medical School Internal Animal Care and Use Committee. Mice were maintained under specific pathogen-free conditions, and food and water were provided ad libitum. C57BL/6 mice were purchased from Charles River and *P48-Cre* strains purchased from Jackson Laboratory. *Trp53^fl/fl^* and *Kras^+/LSL-G12D^* breeding pairs were generously provided by Wen Xue.

### Pancreas transplant tumor models

5×10^4^ *KPC1*, 2.5×10^5^ *KPC2*, 5×10^4^ *KP1*, or 1×10^5^ *KP2* cells were resuspended in 25 μl of Matrigel (Matrigel, BD) diluted 1:1 with cold PBS and transplanted into the pancreas of 8-10 week old C57BL/6 female mice. Following anesthetization using 2-3% isoflurane, an incision was made in the left abdominal side and the cell suspension was injected into the tail region of the pancreas using a Hamilton Syringe. Successful injection was verified by the appearance of a fluid bubble without signs of intraperitoneal leakage. The abdominal wall was sutured with an absorbable Vicryl suture (Ethicon), and the skin was closed with wound clips (CellPoint Scientific Inc.). Mice were monitored for tumor development by ultrasound imaging, and randomized into treatment groups 1-week post-transplantation based on tumor volume. Upon sacrifice pancreas tumor tissue was allocated for 10% formalin fixation, OCT frozen blocks, flow cytometry analysis, and FACS sorting for downstream RNA-seq analysis.

### Lung transplant tumor models

5×10^5^ *KPC1*, 5×10^5^ *KPC2*, 4×10^4^ *KP1*, or 2.5×10^5^ *KP2* cells were resuspended in PBS and transplanted by tail vein injection into 8-10 week old C57BL/6 female mice. Mice were monitored for tumor development by bioluminescence imaging (BLI) on a Xenogen IVIS (Caliper Life Sciences) and randomized into various treatment cohorts 1-week post-transplantation. Upon sacrifice lung lobes were allocated for 10% formalin fixation (1 lobe), OCT frozen blocks (1 lobe), and flow cytometry analysis and FACS sorting (3 lobes).

### Liver transplant tumor models

2×10^5^ *KPC1* or *KP1* cells were resuspended in 25 μl of Matrigel (Matrigel, BD) diluted 1:1 with cold PBS and transplanted directly into the liver of 8-10 week old C57BL/6 female mice. Following tumor development in the liver (6-8 days post-transplantation), mice were evaluated by BLI on a Xenogen IVIS (Caliper Life Sciences) to quantify liver tumor burden before being randomized into various study cohorts. Upon sacrifice liver tumor tissue was allocated for 10% formalin fixation, OCT frozen blocks, and flow cytometry analysis.

### *KPC* GEMM model

*Trp53^fl/+^, Kras^+/LSL-G12D^* and *P48-Cre* strains on a C57Bl/6 background were interbred to obtain *P48-Cre; Kras^+/LSL-G12D^; Trp53^fl/+^* (*KPC*) GEMM mice. Mice were monitored for tumor development by ultrasound imaging, and enrolled and randomized into treatment groups once tumors reached ~50 mm^3^ in volume. Upon sacrifice pancreas tumor tissue was allocated for 10% formalin fixation and OCT frozen blocks.

### Preclinical drug studies

Mice were treated with vehicle, trametinib (1 mg/kg body weight), palbociclib (100 mg/kg body weight) and/or tazemetostat (125 mg/kg (low) or 400 mg/kg (high) body weight) *per os* for 4 consecutive days followed by 3 days off treatment. For NK and T cell depletion, mice were injected intraperitoneally (IP) with an αNK1.1 (250 μg; PK136, BioXcell), αCD8 (200 μg; 2.43, BioXcell) or αCD4 (200 μg; GK1.5, BioXcell) antibody twice per week. Depletion of NK, CD4^+^, and CD8^+^ T cells was confirmed by flow cytometric analysis. For neutralization of chemokine signaling, mice were injected IP with an αCCL2 (200 μg; 2H5, BioXcell) or αCXCR3 (200 μg; CXCR3-173, BioXcell) antibody twice per week. No obvious toxicities were observed in treated animals. Ultrasound imaging was repeated every 2 weeks during treatment to assess changes in PDAC tumor burden.

### Ultrasound Imaging

High-contrast ultrasound imaging was performed on a Vevo 3100 System with a MS250 13- to 24-MHz scanhead (VisualSonics) to stage and quantify PDAC tumor burden. Tumor volume was analyzed using Vevo 3100 software.

### Bioluminescence imaging

Bioluminescence imaging (BLI) was used to track luciferase expression in transplanted *KPC1* PDAC and *KP1* LUAD tumor cells expressing a luciferase-GFP reporter. Mice were injected IP with luciferin (5 mg/mouse; Gold Technologies) and then imaged on a Xenogen IVIS Spectrum imager (PerkinElmer) 10-15 minutes later for 60 seconds. Quantification of luciferase signaling was analyzed using Living Image software (Caliper Life Sciences).

### Flow cytometry

For analysis of MHC-I expression in cell lines cultured *in vitro*, cells were treated for 8 days with vehicle (DMSO), combined trametinib (25 nM) and palbociclib (500 nM), and/or tazmetostat (5 μM) and then trypsinized, resuspended in PBS supplemented with 2% FBS, and stained with an H-2k^b^ antibody (AF6-88.5.5.3, eBioscience) for 30 minutes on ice. Flow cytometry was performed on a BD LSR II, and data were analyzed using FlowJo (TreeStar).

For *in vivo* sample preparation, lungs were isolated, flushed with PBS, and allocated for 10% formalin fixation (1 lobe), OCT frozen blocks (1 lobe), and FACS (3 lobes) following 2-week treatment. Pancreatic tumor tissue was isolated from the spleen and normal tissue and allocated for 10% formalin fixation, OCT frozen blocks, and FACS following 2-week treatment. Liver tumors were isolated from liver lobes and allocated for 10% formalin fixation, OCT frozen blocks, and FACS following 2-week treatment. To prepare single cell suspensions for flow cytometry analysis, lung, pancreas, or liver tissue was minced with scissors into small pieces and placed in 5ml of collagenase buffer (1x HBSS w/ calcium and magnesium (Gibco), 1 mg/ml Collagenase A (Roche) for LUAD tumors or Collagenase V (Sigma-Aldrich) for PDAC tumors, and 0.1 mg/ml DNase I) in C tubes and then processed using program 37C_m_LDK_1 (for LUAD tumors) or 37C_m_TDK1_1 (for PDAC tumors) on a gentleMACS Octo dissociator with heaters (Miltenyi Biotec). Spleens were placed in 3 ml of PBS supplemented with 2% FBS in C tubes and dissociated using program m_spleen_01 on a gentleMACS Octo dissociator with heaters (Miltenyi Biotec). Dissociated tissue was passaged through a 70 μm cell strainer and centrifuged at 1500 rpm x 5 minutes. Red blood cells were lysed with ACK lysis buffer (Quality Biological) for 5 minutes, and samples were centrifuged and resuspended in PBS supplemented with 2% FBS. Samples were blocked with anti-CD16/32 (FC block, BD Pharmigen) for 20 minutes and then incubated with the following antibodies for 30 minutes on ice: CD45 (30-F11), NK1.1 (PK136), CD3 (17A2), CD8 (53-6.7), CD4 (GK1.5), CD69 (H1.2F3), Sca-1 (Ly6A/E; D7), F4/80 (BM8) (Biolegend); and CD11b (M1/70) BD Biosciences). NK cells were gated from the CD45^+^CD3^−^NK1.1^+^ population. DAPI was used to distinguish live/dead cells, and tumor cells were gated as GFP^+^. Flow cytometry was performed on an BD LSRFortessa or LSR II, and data were analyzed using FlowJo (TreeStar).

For analysis of Granzyme B (GZMB) expression in NK and T cells, single cell suspensions from tumor tissue were resuspended in RPMI media supplemented with 10% FBS and 100 IU/ml P/S and incubated for 4 hours with PMA (20 ng/ml, Sigma-Aldrich), Ionomycin (1 μg/ml, STEMCELL technologies), and monensin (2 μM, Biolegend) in a humidified incubator at 37°C with 5% CO_2_. Cell surface staining was first performed with CD45 (30-F11), NK1.1 (PK136), CD3 (17A2), CD8 (53-6.7), and CD4 (GK1.5) antibodies (Biolegend). Intracellular staining was then performed using the Foxp3/transcription factor staining buffer set (eBioscience), where cells were fixed, permeabilized, and then stained with a GZMB antibody (GB11; Biolegend). GZMB expression was evaluated by gating on CD3^−^NK1.1^+^ NK cells and CD3^+^CD8^+^ T cells on an BD LSR II flow cytometer as described above.

### NK and T cell degranulation assays

Mice were injected intravenously (i.v.) with 250 μl of a solution containing 25 μg anti-CD107a PE (ID4B, Biolegend) and 10 μg monensin (Biolegend) in PBS 4 hours before mice were euthanized. Tumor tissue was then isolated, dissociated into single cell suspensions, stained with cell surface antibodies, and analyzed by flow cytometry as described above.

### Immunohistochemistry (IHC)

Tissues were fixed overnight in 10% formalin, embedded in paraffin, and cut into 5 μm sections. Haematoxylin and eosin (H&E), Masson’s trichrome, and immunohistochemical staining were performed using standard protocols. Sections were de-paraffinized, rehydrated, and boiled in a pressure cooker for 20 minutes in 10 mM citrate buffer (pH 6.0) for antigen retrieval. Antibodies were incubated overnight at 4°C. The following primary antibodies were used: pERK^T202/Y204^ (4370), EZH2 (5246) (Cell Signaling); Ki67 (AB16667), H3K27me3 (AB177178), CD3 (AB5690), GZMB (AB4059), CD31 (AB28364) (Abcam); pRB^S807/S811^ (Sc-16670, Santa Cruz); and NKp46 (AF2225, R&D Systems). HRP-conjugated secondary antibodies (Vectastain Elite ABC Kits: PK-6200; Rabbit, PK-6101; Goat, PK-6105) were applied for 30 minutes and visualized with DAB (Vector Laboratories; SK-4100). For quantification of CD31^+^ blood vessels and NKp46^+^, CD3^+^, and GZMB^+^ immune cells, 5-10 high power 20x fields per section were counted and averaged.

### High throughput RNA-sequencing (RNA-seq)

For RNA-seq analysis of PIP, PIL, LIL, and LIP tumor samples, GFP^+^ tumor cells were FACS sorted on a FACSAria (BD Biosciences) from the lungs or pancreas of tumor-bearing mice following 2-week treatment with vehicle or combined trametinib (1 mg/kg body weight) and palbociclib (100 mg/kg). Total RNA was extracted from tumor cells using the RNeasy Mini Kit (Qiagen). Purified polyA mRNA was subsequently fragmented, and first and second strand cDNA synthesis performed using standard Illumina mRNA TruSeq library preparation protocols. Double stranded cDNA was subsequently processed for TruSeq dual-index Illumina library generation. For sequencing, pooled multiplexed libraries were run on a HiSeq 2500 machine on RAPID mode. Approximately 10 million 76bp single-end reads were retrieved per replicate condition. RNA-Seq data was analyzed by removing adaptor sequences using Trimmomatic^85^, aligning sequencing data to GRCh37.75(hg19) with STAR^86^, and genome wide transcript counting using HTSeq^87^ to generate a RPKM matrix of transcript counts. Genes were identified as differentially expressed using R package DESeq2 with a cutoff of absolute log_2_FoldChange ≥ 1 and adjusted p-value <0.05 between experimental conditions^78^. For heatmap visualization of selected genes and pathways, samples were z-score normalized and plotted using ‘pheatmap’ package in R. Heatmaps display the fold change in RPKM expression values from T/P-treated samples normalized to vehicle-treated samples (T/P-V). Over-representation analysis of DEGs against KEGG^88^ Pathways was performed using clusterProfiler^83^.

For RNA-seq analysis of *shRen* and *shEzh2* PDAC tumor samples, GFP^+^ tumor cells were FACS sorted on a FACSAria (BD Biosciences) from the pancreas of tumor-bearing mice following 2-week treatment with vehicle or combined trametinib (1 mg/kg body weight) and palbociclib (100 mg/kg). Total RNA was extracted from tumor cells using the RNeasy Mini Kit (Qiagen). Library preparation and sequencing on a NovaSeq 6000 was performed by Novogene. Approximately 30 million 150bp paired-end reads were retrieved per replicate condition. Quality of raw sequencing data was checked using FastQC (https://www.bioinformatics.babraham.ac.uk/projects/fastqc/) to assure data quality. Paired-end reads were aligned to the mouse reference genome GRCm38 (Ensembl, version 101) using STAR^86^. A gene-by-sample count matrix was generated using featureCounts^89^. All downstream statistical analyses were done using the R programming language^90^. Briefly, entries of genes with extremely low expression were first removed from the gene-by-sample count matrix. Differential gene analysis was performed using DESeq2^78^ in consideration of surrogate variables for hidden variations, which were identified using svaseq^79^. Genes with absolute values of log_2_ (fold change) greater than one and *p*-values less than 0.05, which were corrected for multiple testing using the Benjamini-Hochberg procedure^80^, were considered as significantly differentially expressed genes (DEGs). Over-representation analysis of DEGs against KEGG Pathways^88^ was performed using clusterProfiler^83^. Heatmaps were generated using pheatmap (https://github.com/raivokolde/pheatmap).

To assess expression of SASP genes in human PDAC and LUAD cell lines treated with vehicle (DMSO), trametinib (25 nM), and/or palbociclib (500 nM) for 8 days in culture, we interrogated a previously published RNA-seq dataset under the GEO accession number GSE110397^19^. Graphs display normalized RPKM expression values.

### Gene Set Enrichment Analysis (GSEA)

GSEA was performed using the GSEAPreranked tool for conducting gene set enrichment analysis of data derived from RNA-seq experiments (version 2.07) against Hallmark signatures in the MSigDB database (http://software.broadinstitute.org/gsea/msigdb) and published senescence signatures^40^. The metric scores were calculated using the sign of the fold change multiplied by the inverse of the p-value.

### Transcription factor enrichment analysis

Transcription factor enrichment analysis was performed using gene set libraries from Enrichr^91^. Significance of the tests was assessed using combined score, described as c = log(p) * z, where c is the combined score, p is Fisher exact test p-value, and z is z-score for deviation from expected rank.

### Pearson’s correlation analysis

Gene expression data from 145 primary PDAC tumors (GSE71729)^58^ was downloaded with GEOquery2 package. Correlation analysis between PRC2^92^ and our custom EZH2 repression signatures (generated from list of genes downregulated in *shEzh2* compared to *shRen* PDAC tumor cells from RNA-seq analysis in Fig. 5), inflammatory response, NK cell^93^, and CD8^+^ T cell^94^ gene sets (see Supplementary Table 8), and *CCL2*, *CXCL9*, and *CXCL10* expression was performed using ggpubr package. Pearson’s correlation coefficient (R) values are displayed.

### Human PDAC specimens

PDAC patient samples were derived from surgical candidates undergoing Whipple procedures consented under the IRB approved protocol no. H-4721. De-identified FFPE tumor specimens were cut into 5 μm sections and IHC performed as described above to stain for human EZH2 (5246, Cell Signaling) and NKp46 (AF1850, R&D Systems). EZH2 staining was scored as high (strong nuclear staining throughout tumor), intermediate (nuclear staining in some but not all tumor areas), or low (little to no positive staining in the tumor). NKp46^+^ NK cell numbers were scored as high (> 5 cells per 40x field), intermediate (2-4 cells per 40x field), or low (< 2 cells per 40x field). Survival data from some PDAC patients was also available through the IRB approved protocol no. H-4721.

### Statistical analysis

Statistical analyses were performed as described in the figure legend for each experiment. Data are expressed as mean ± s.e.m. Group size was determined on the basis of the results of preliminary experiments and no statistical method was used to predetermine sample size. The indicated sample size (*n*) represents biological replicates. Group allocation and outcome assessment were not performed in a blinded manner. All samples that met proper experimental conditions were included in the analysis. All experiments were repeated independently 2-3 times. Survival was measured using the Kaplan–Meier method. Statistical significance was determined by Student’s *t*-test and log-rank test using Prism 6 software (GraphPad Software) as indicated. Significance was set at *P* < 0.05.

## DATA AVAILABILITY

RNA-seq and CUT&Tag data have been deposited in the Gene Expression Omnibus (GEO) under accession nos. GSE201495, GSE141684, and GSE203623. Gene expression data for human LUAD and PDAC cell lines treated with T/P were obtained under accession no. GSE110397. Gene expression data from 145 primary human PDAC specimens were obtained under accession no. GSE71729. Custom gene sets are provided in Supplementary Table 8. Any code used in this manuscript will be made available upon request. Further information and requests for resources and reagents should be directed to the corresponding author, M.R.

## ACKNOWLEDGEMENTS

We thank K. Hatzi for providing shRNA constructs; R.Mezzadra for generating *Ccl2* O/E cell lines; G. Cottle for technical assistance; and Wen Xue, Art Mercurio, Michelle Kelliher, Michael Green, Jane Chuprin, Lin Zhou, and other members of the Ruscetti laboratory for helpful suggestions and comments on the manuscript. Some figures were created with Biorender.com. This work was supported by a K99/R00 CA241110 grant from the National Cancer Institute (NCI) to M.R and a Memorial Sloan Kettering Cancer Center Support grant (P30 CA008748) to S.W.L.. We acknowledge support from Our Danny Postdoc Fund (to L.C.), the NIH (R01 HD072122 to T.G.F, P30 CA008748 S5 to E.d.S.), and the National Center for Advancing Translational Sciences (UL1-TR001453 to K.S.). S.W.L. is the Geoffrey Beene Chair for Cancer Biology and a Howard Hughes Medical Institute investigator.

## AUTHOR CONTRIBUTIONS

L.C. and M.R. conceived the study, designed and performed experiments, interpreted results, and wrote the paper with assistance from all authors. K.C.M., Y.L.-D., K.D.D., and J.P.M. designed, performed and analyzed *in vitro* and *in vivo* experiments. K.C.M., Y.L.-D., K.D.D., J.S., W.L., A.K., and E.d.S. produced and treated animal models. H.L., J.L., Y.-j.H., and L.J.Z. analyzed transcriptomic datasets. S.G., T.G.F., H.L., and L.J.Z. designed and performed CUT&Tag analysis. M.F. and K.S. provided human PDAC patient specimens and data. M.R. and S.W.L. supervised the study.

---

**Extended Data Fig. 1.**
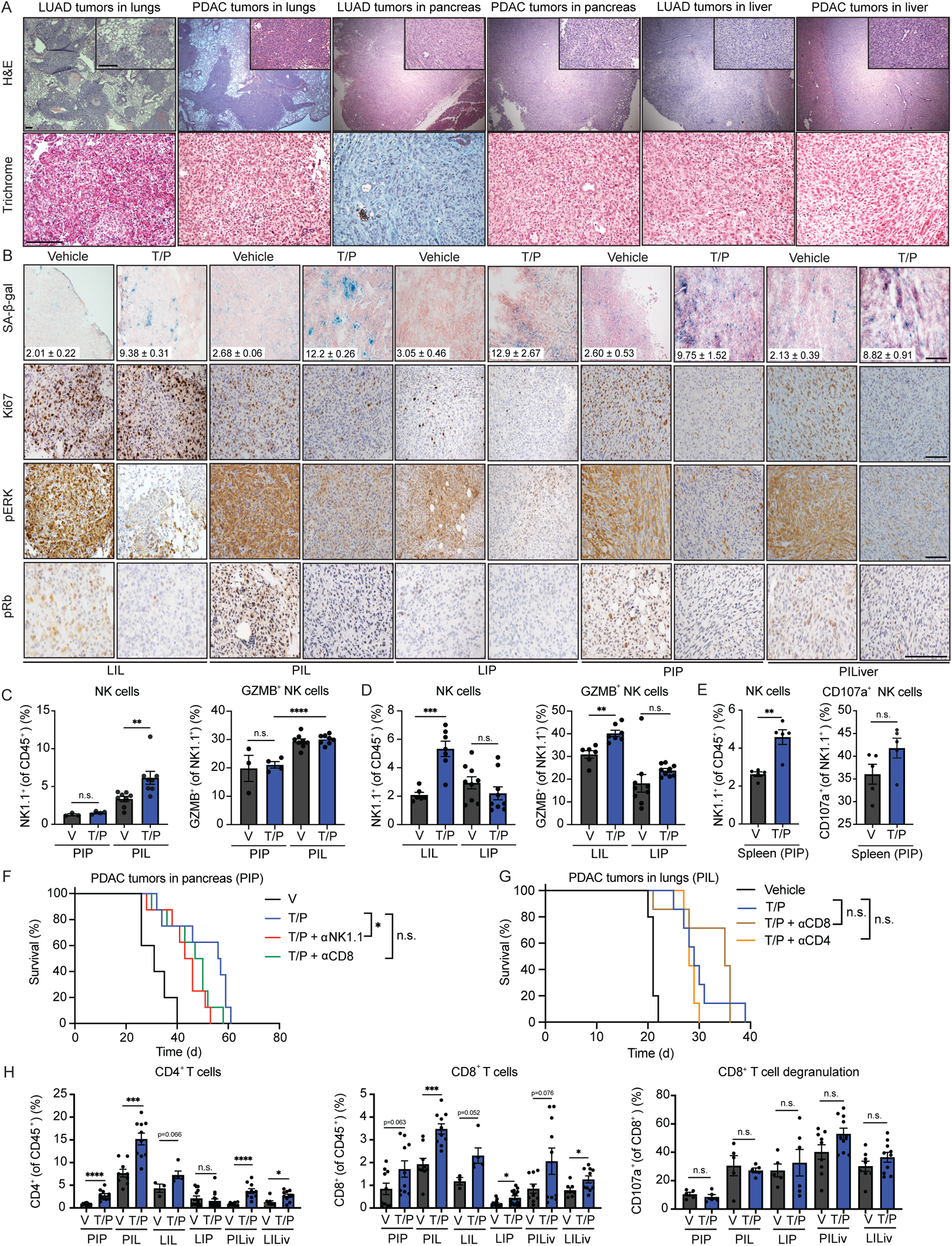
Suppression of NK cell immunity specific to pancreas TME despite similar tumor, senescence, and T cell responses to T/P treatment across tumor conditions. **a**, Haematoxylin and eosin (H&E) (top) and Masson’s trichrome (bottom) staining of indicated *KPC1* PDAC (PIP, PIL, PILiver) and *KP1* LUAD (LIL, LIP, LILiver) derived-tumors. Scale bars, 100μm. **b**, Immunohistochemical (IHC) staining of indicated *KPC1* PDAC (PIP, PIL, PILiver) and *KP1* LUAD (LIL, LIP) derived-tumors treated with vehicle (V) or combined trametinib (1mg/kg) and palbociclib (100 mg/kg) (T/P) for 2 weeks. Quantification of SA-β-gal positive area is shown (n=2-3 per group; T/P versus V (LIL), p < 0.0001; T/P versus V (PIL), p < 0.0001; T/P versus V (LIP), p = 0.066; T/P versus V (PIP), p = 0.047; T/P versus V (PILiv), p = 0.0025). Scale bars, 100μm. **c-d**, *KPC2* PDAC or *KP2* LUAD tumor cells expressing GFP were injected i.v. or orthotopically into the pancreas of C57BL/6 mice. Following tumor formation in the lungs or pancreas, mice were treated as in (**b**). Flow cytometry analysis of NK cell numbers and degranulation in PDAC (PIP, PIL) (**c**) and LUAD-derived tumors (LIL, LIP) (**d**) grown in different organs are shown (n ≥ 3 per group). **e**, Flow cytometry analysis of NK cell numbers and degranulation in the spleens of mice with *KPC1* PDAC tumors orthotopically transplanted into the pancreas (PIP) treated as in (**b**) (n=5 per group). **f**, Kaplan-Meier survival curve of mice with *KPC2* PDAC tumors orthotopically transplanted into the pancreas (PIP) treated with vehicle, combined trametinib (1mg/kg) and palbociclib (100mg/kg), and/or depleting antibodies against NK1.1 (PK136; 250 μg) or CD8 (2.43; 200 μg) (n ≥ 5 per group). **g**, Kaplan-Meier survival curve of mice with *KPC1* PDAC tumors i.v. injected into the lungs (PIL) treated with vehicle, combined trametinib (1mg/kg) and palbociclib (100mg/kg), and/or depleting antibodies against CD8 (2.43; 200 μg) or CD4 (GK1.5; 200 μg) (n ≥ 5 per group). **h**, Flow cytometry analysis of CD4^+^ and CD8^+^ T cell numbers and degranulation in *KPC1* PDAC (PIP, PIL, PILiver) and *KP1* LUAD-derived tumors (LIL, LIP, LILiver) grown in different organs and treated as in (**b**) (n ≥3 per group). *P* values in **c-e** and **h** were calculated using two-tailed, unpaired Student’s t-test, and those in **f** and **g** were calculated using log-rank test. Error bars, mean ± SEM. **** *P* <0.0001, *** *P* <0.001, ** *P* <0.01, * *P* <0.05. n.s., not significant.

**Extended Data Fig. 2.**
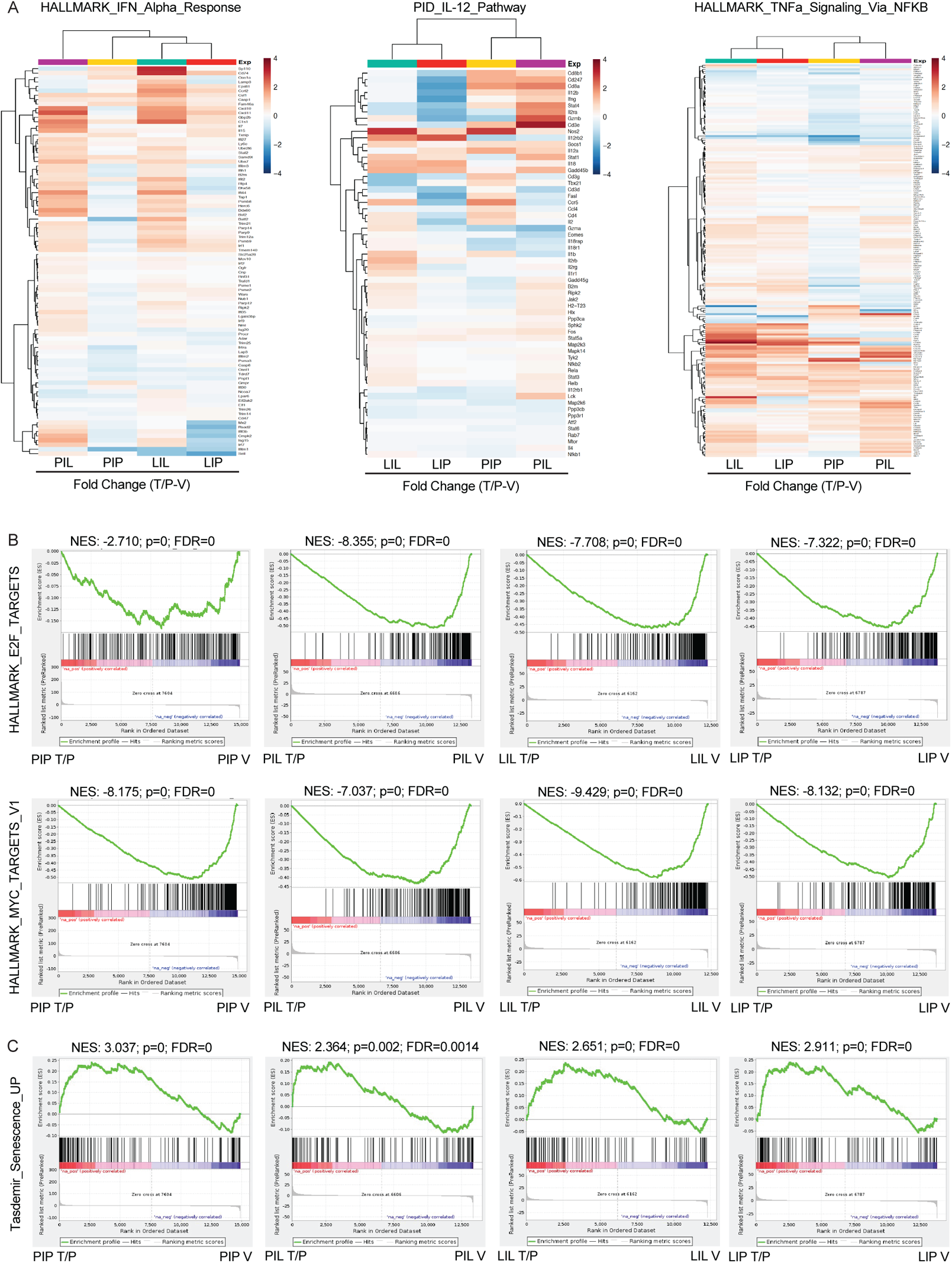
Pancreatic tumors undergo senescence but have an altered pro-inflammatory SASP following T/P treatment. **a**, Heatmaps showing fold change in IFNα (left), IL-12 (middle), and TNFα pathway genes (right) following T/P treatment in indicated tumor settings from RNA-seq data in Figure 2a (n=2-4 per group). **b-c**, Gene Set Enrichment Analysis (GSEA) of RNA-seq data in Fig. 2a (n=2-4 per group). NES, normalized enrichment score.

**Extended Data Fig. 3.**
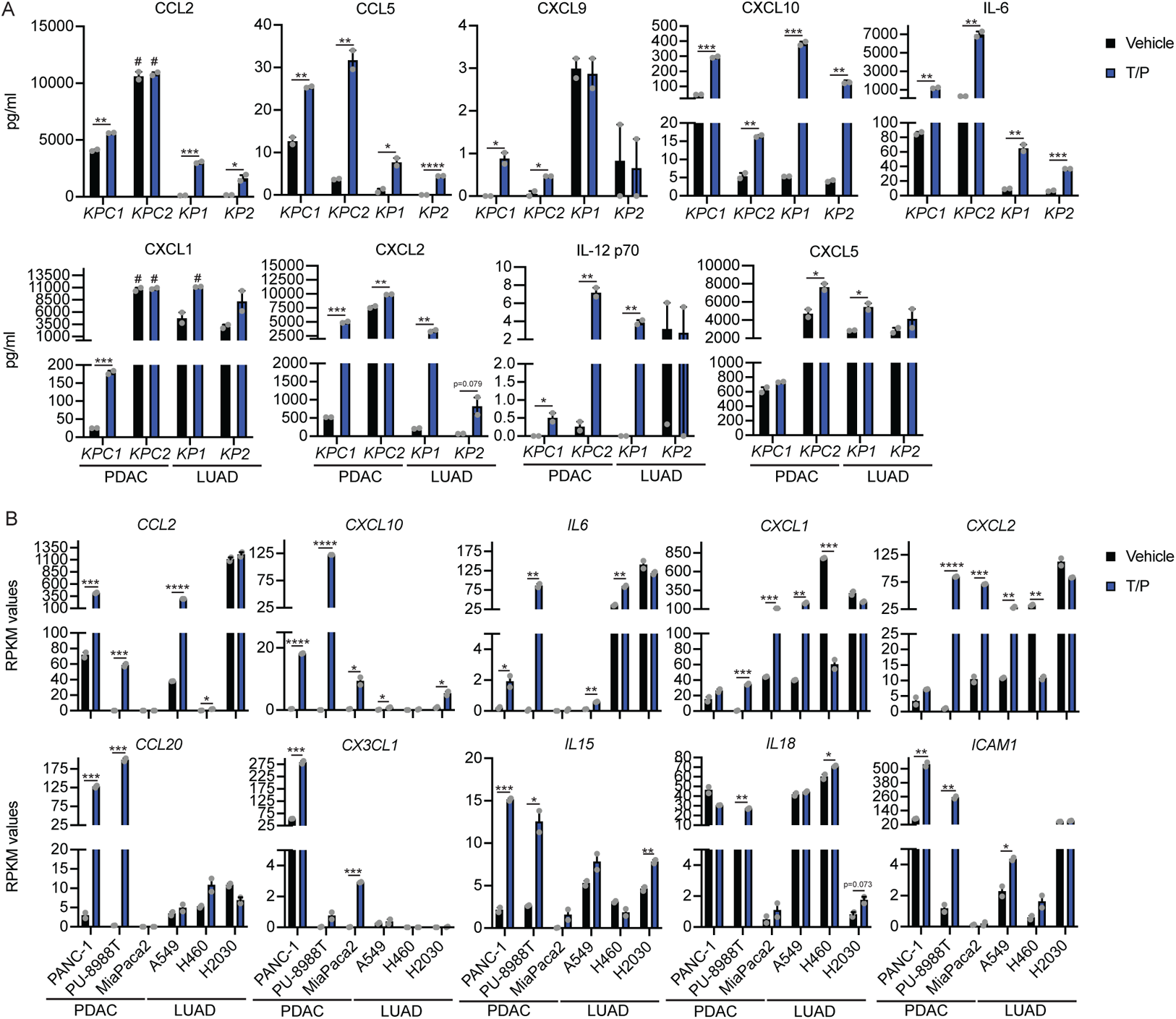
PDAC and LUAD tumor cells have a similar pro-inflammatory SASP response to T/P-induced senescence *in vitro*. **a**, Cytokine array analysis of pro-inflammatory SASP genes in murine PDAC and LUAD cell lines treated with vehicle or combined trametinib (25 nM) and palbociclib (500nM) for 8 days (n=2). #, outside the detectable limit. **b**, Normalized expression levels of pro-inflammatory SASP genes in human PDAC and LUAD cell lines following treatment as in (**a**) from analysis of RNA-seq data generated in Ruscetti et al. (2018)^19^ (n=2 per group). *P* values in **a** and **b** were calculated using two-tailed, unpaired Student’s t-test. Error bars, mean ± SEM. **** *P* <0.0001, *** *P* <0.001, ** *P* <0.01, * *P* <0.05. n.s., not significant.

**Extended Data Fig. 4.**
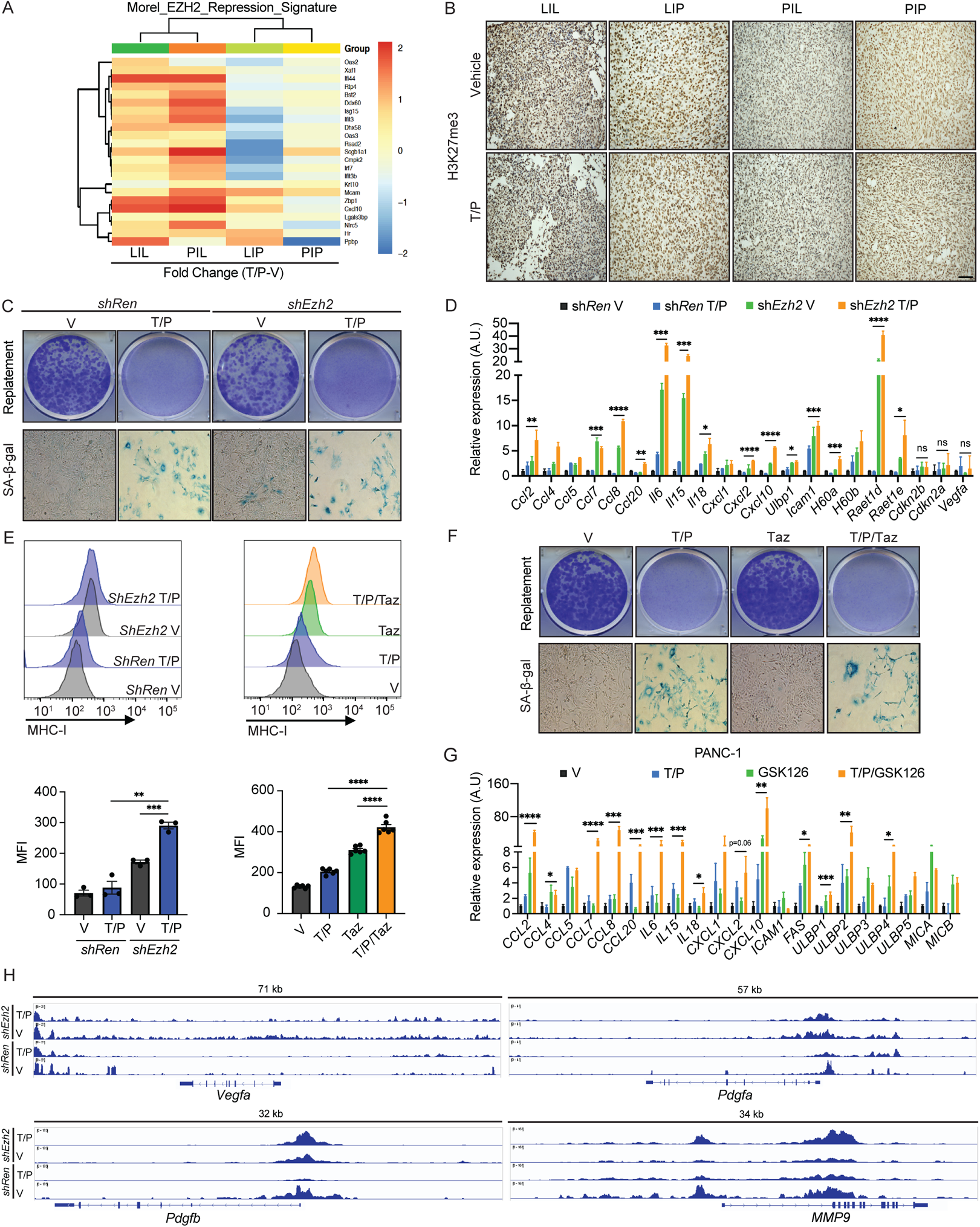
Suppression of EZH2 activity in PDAC induces pro-inflammatory SASP and immunomodulatory cell surface molecules following T/P treatment without impacting senescence-associated cell cycle arrest or pro-angiogenic SASP factors. **a**, Heatmap showing fold change in EZH2 repressed genes^95^ following T/P treatment from RNA-seq data in Fig. 2a (n=2-4 per group). **b**, IHC staining of indicated *KPC1* PDAC (PIP, PIL) and *KP1* LUAD (LIL, LIP) derived-tumors grown in different organs and treated with vehicle (V) or combined trametinib (1mg/kg) and palbociclib (100 mg/kg) (T/P) for 2 weeks. Scale bars, 50μm. **c**, Representative clonogenic assay images (from 3 biological replicates) of *KPC1* PDAC cells harboring *shRen* or *shEzh2* shRNAs replated in the absence of drugs after an 8-day pre-treatment with vehicle or combined trametinib (25 nM) and palbociclib (500 nM) (top). Bottom, representative SA-β-gal staining (from 3 biological replicates) of *KPC1* PDAC cells harboring *shRen* or *shEzh2* shRNAs and treated with vehicle or combined trametinib (25 nM) and palbociclib (500 nM) for 8 days. **d**, qRT-PCR analysis of senescence and SASP gene expression in *KPC1* PDAC cells harboring *shRen* or *shEzh2* shRNAs treated with vehicle or combined trametinib (25 nM) and palbociclib (500 nM) for 8 days (n=3). A.U., arbitrary units. **e**, Representative histograms (top) and quantification of mean fluorescent intensity (MFI) of MHC-I (H-2k^b^) expression (bottom) on *KPC1* PDAC cells harboring *shRen* or *shEzh2* shRNAs (left) or parental *KPC1* PDAC cells (right) treated with vehicle, combined trametinib (25 nM) and palbociclib (500nM), and/or tazemetostat (5 μM) for 8 days (n=3-6). **f**, Representative clonogenic assay images (from 3 biological replicates) of *KPC1* PDAC cells replated in the absence of drugs after an 8-day pre-treatment with vehicle, combined trametinib (25 nM) and palbociclib (500 nM), and/or tazemetostat (5 μM) (top). Bottom, representative SA-β-gal staining (from 3 biological replicates) of *KPC1* PDAC cells treated with vehicle, combined trametinib (25 nM) and palbociclib (500 nM), and/or tazemetostat (5 μM) for 8 days. **g**, qRT-PCR analysis of SASP gene expression in human PANC-1 PDAC cells treated with vehicle, trametinib (25 nM), palbociclib (500 nM), and/or GSK126 (1 μM) for 8 days (n=3). A.U., arbitrary units. **h**, Genome browser tracks from CUT&Tag analysis showing H3K27me3 occupancy at select pro-angiogenic SASP gene loci in *KPC1* PDAC cells harboring *Ren* or *Ezh2* shRNAs treated with vehicle or trametinib (25 nM) and palbociclib (500 nM) for 8 days (n=2-4 per group). *P* values in **d**, **e** and **g** were calculated using two-tailed, unpaired Student’s t-test. Error bars, mean ± SEM. **** *P* <0.0001, *** *P* <0.001, ** *P* <0.01, * *P* <0.05. n.s., not significant.

**Extended Data Fig. 5.**
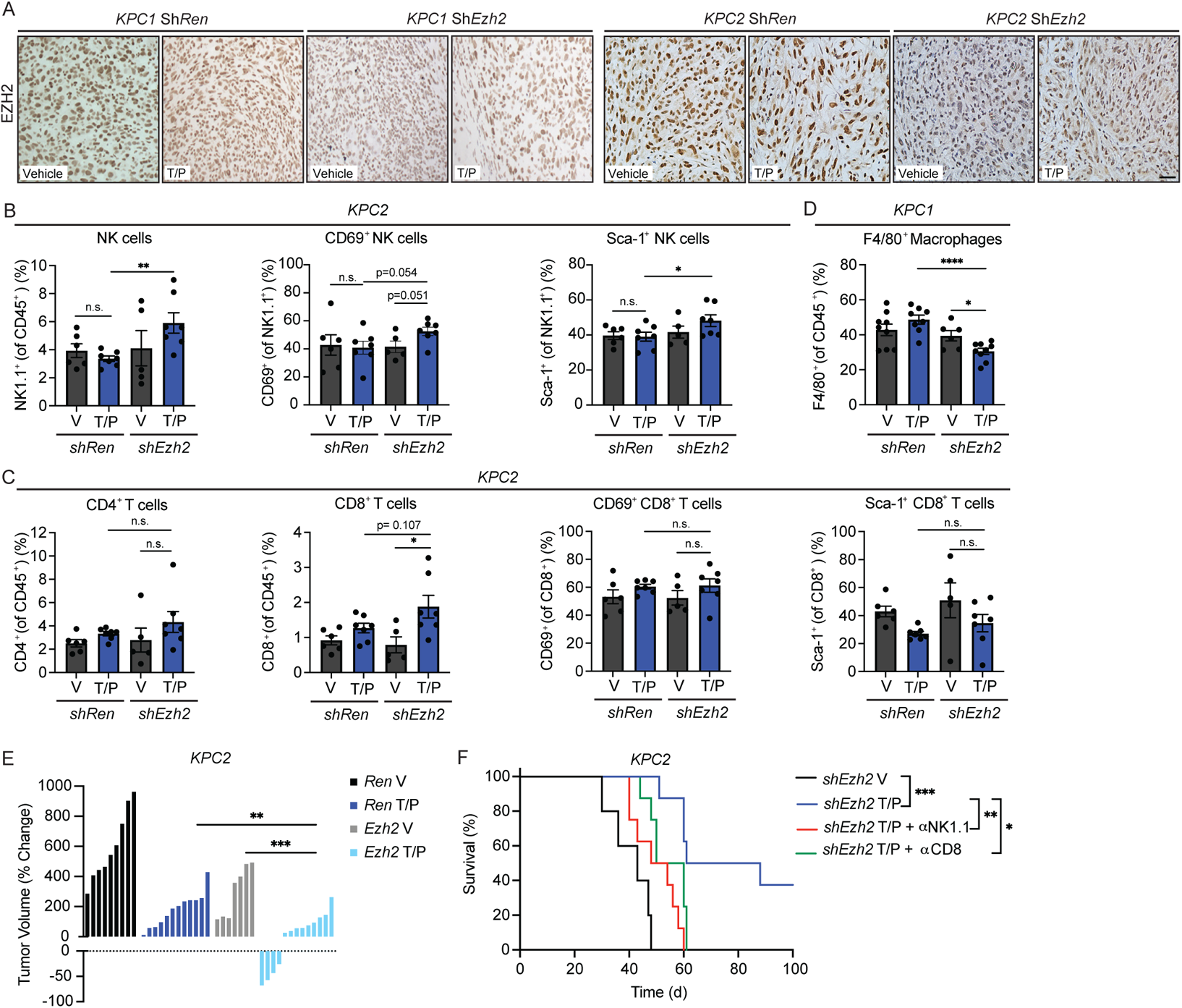
EZH2 knockdown in the *KPC2* PDAC orthotopic transplant model also potentiates anti-tumor NK and CD8^+^ T cell immunity and long-term tumor regressions following T/P treatment. **a**, IHC staining of *KPC1* (left) or *KPC2* (right) orthotopic PDAC tumors harboring *shRen* or *shEzh2* shRNAs treated with vehicle or combined trametinib (1mg/kg) and palbociclib (100 mg/kg) (T/P) for 2 weeks. Scale bars, 50μm. **b-c**, Flow cytometry analysis of NK cell (**b**) and T cell (**c**) numbers and activation markers in *KPC2* orthotopic PDAC tumors harboring indicated shRNAs treated as in (**a**) (n ≥ 5 per group). **d**, Flow cytometry analysis of F4/80^+^ macrophages in *KPC1* orthotopic PDAC tumors harboring indicated shRNAs treated as in (**a**) (n ≥ 6 per group). **e**, Waterfall plot of the response of *KPC2* orthotopic PDAC tumors harboring indicated shRNAs to treatment as in (**a**) (n ≥ 7 per group). **f**, Kaplan-Meier survival curve of mice with *shEzh2 KPC2* orthotopic PDAC tumors treated with vehicle, combined trametinib (1mg/kg) and palbociclib (100mg/kg), and/or depleting antibodies against NK1.1 (PK136; 250 μg) or CD8 (2.43; 200 μg) (n ≥ 5 per group). *P* values in **b-e** were calculated using two-tailed, unpaired Student’s t-test, and those in **f** were calculated using log-rank test. Error bars, mean ± SEM. **** *P* <0.0001, *** *P* <0.001, ** *P* <0.01, * *P* <0.05. n.s., not significant.

**Extended Data Fig. 6.**
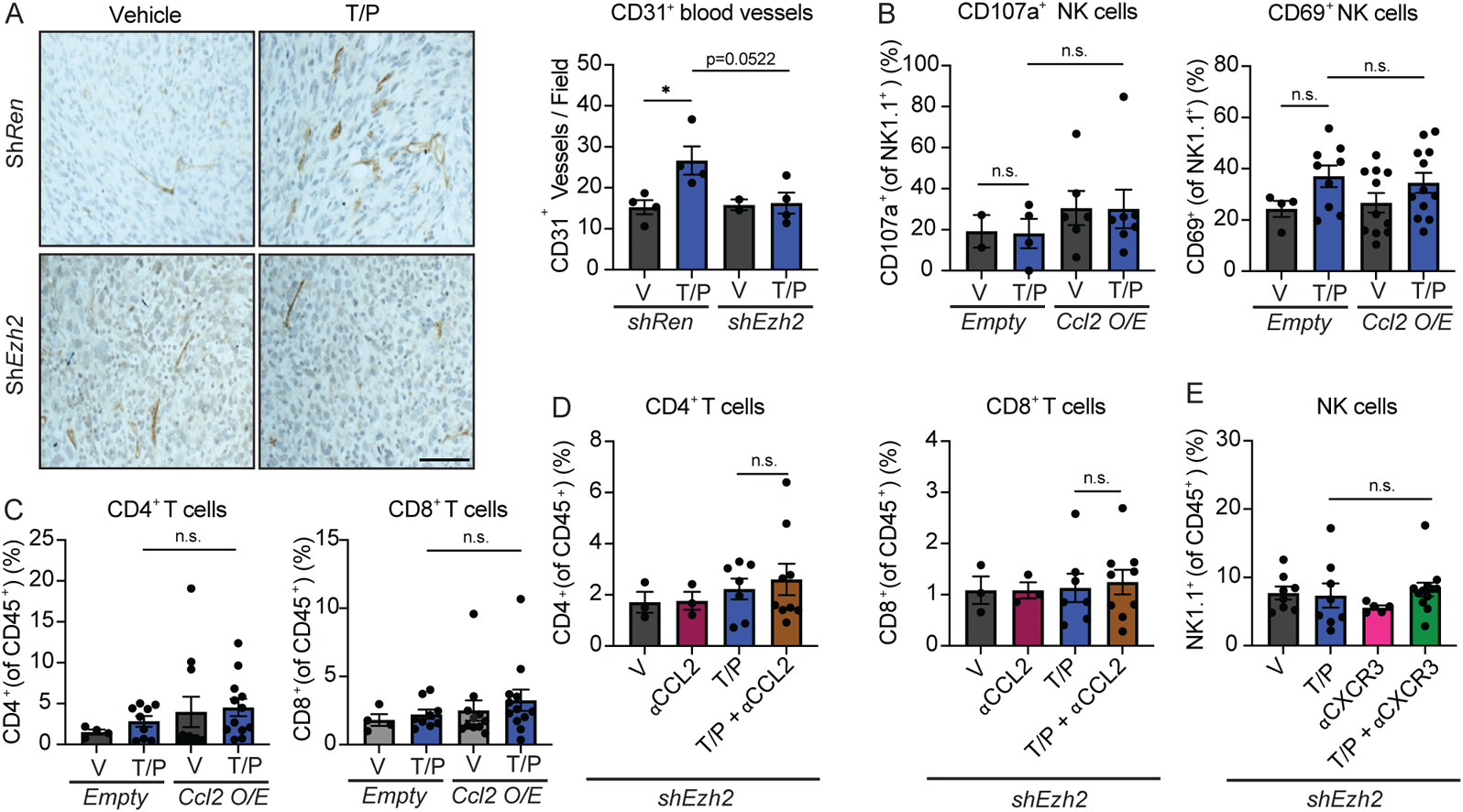
EZH2 blockade reduces T/P-induced blood vessel formation and promotes CCL2 and CXCL9/10 secretion that increases NK and CD8^+^ T cell infiltration into PDAC. **a**, IHC staining of *KPC1* orthotopic PDAC tumors harboring *shRen* or *shEzh2* shRNAs treated with vehicle or combined trametinib (1mg/kg) and palbociclib (100 mg/kg) (T/P) for 2 weeks. Scale bars, 50μm. Quantification of blood vessels per field are shown on right (n=2-4). **b-c**, Flow cytometry analysis of NK cell activation markers (**b**) and CD4^+^ and CD8^+^ T cell numbers (**c**) in *KPC1* orthotopic PDAC tumors expressing control *Empty* or *Ccl2* vectors and treated as in (**a**). (n ≥ 2 per group). d, Flow cytometry analysis of CD4^+^ and CD8^+^ T cell numbers in *shEzh2 KPC1* orthotopic PDAC tumors following treatment with vehicle, combined trametinib (1mg/kg) and palbociclib (100mg/kg), and/or a CCL2 depleting antibody (2H5; 200 μg) for 2 weeks (n ≥ 3 per group). **e**, Flow cytometry analysis of NK cell numbers in *shEzh2 KPC1* orthotopic PDAC tumors following treatment with vehicle, combined trametinib (1mg/kg) and palbociclib (100mg/kg), and/or a CXCR3 depleting antibody (CXCR3-173; 200 μg) for 2 weeks (n ≥ 5 per group). *P* values in **a-e** were calculated using two-tailed, unpaired Student’s t-test. Error bars, mean ± SEM. **** *P* <0.0001, *** *P* <0.001, ** *P* <0.01, * *P* <0.05. n.s., not significant.

## SUPPLEMENTAL FIGURE TITLES

**Supplementary Fig. 1.** FACS gating strategy

## SUPPLEMENTAL TABLE TITLES

**Supplementary Table 1. Normalized expression counts for LIL, LIP, PIL, and PIP samples from RNA-seq analysis.**

**Supplementary Table 2. Differentially expressed H3K27me3 peaks in *shEzh2* T/P vs. *shRen* T/P samples from CUT&Tag analysis.**

**Supplementary Table 3. Differentially expressed H3K27me3 peaks in *shRen* T/P vs. *shRen* V samples from CUT&Tag analysis.**

**Supplementary Table 4. Differentially expressed H3K27me3 peaks in *shEzh2* T/P vs. *shEzh2* V samples from CUT&Tag analysis.**

**Supplementary Table 5. Differentially expressed H3K27me3 peaks in *shEzh2* V vs. *shRen* V samples from CUT&Tag analysis.**

**Supplementary Table 6. Differentially expressed H3K27me3 peaks in *shEzh2* T/P vs. *shRen* V samples from CUT&Tag analysis.**

**Supplementary Table 7. Normalized expression counts for *shRen* and *shEzh2* PDAC samples from RNA-seq analysis.**

**Supplementary Table 8. Custom gene sets. Supplementary Table 9. qRT-PCR primer sequences.**

