## Supplementary figures and images for "EZH2 inhibition remodels the inflammatory senescence-associated secretory phenotype to potentiate pancreatic cancer immune surveillance"

### Supplementary Figure 1

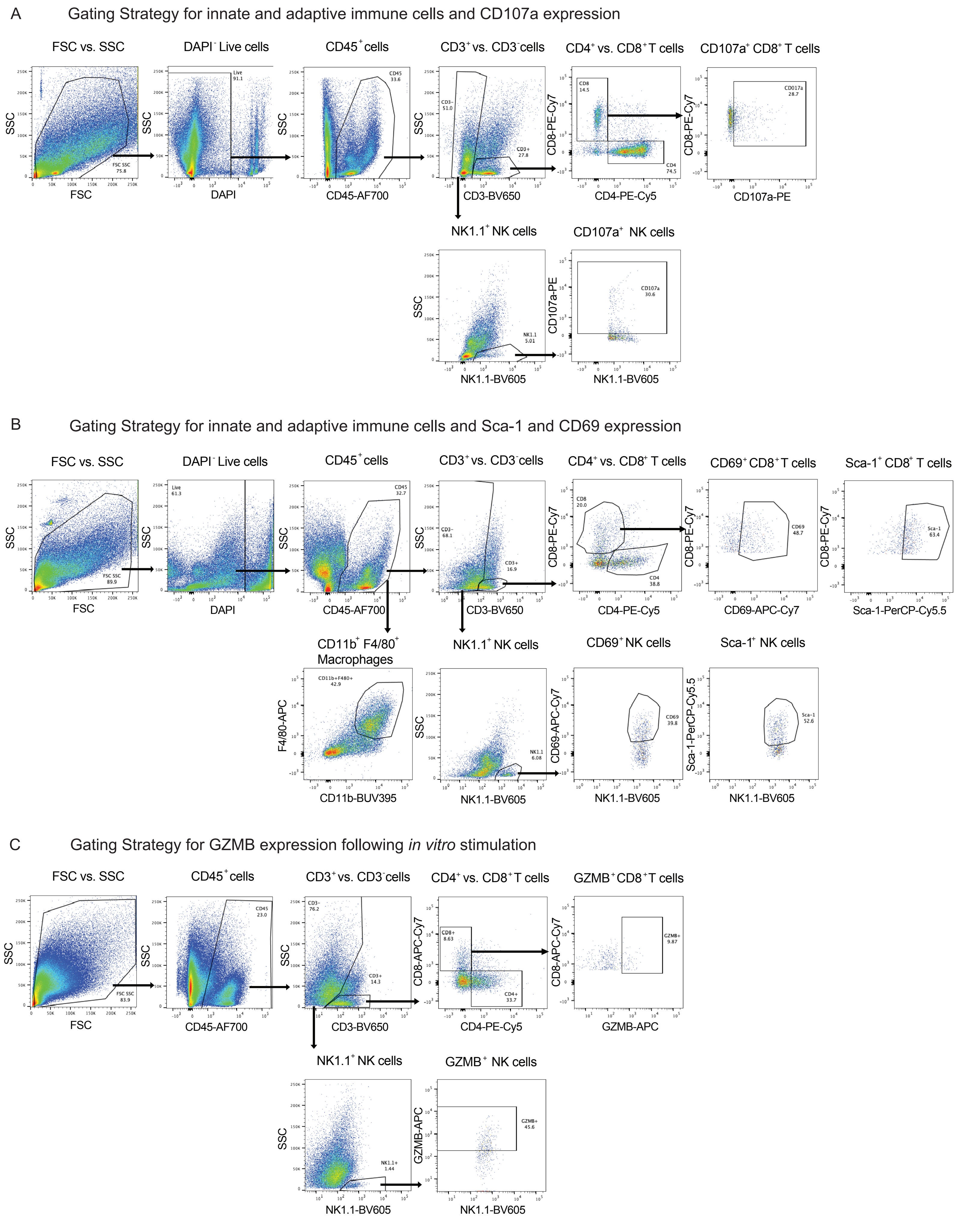
